# Codon affinity in mitochondrial DNA shapes evolutionary and somatic fitness

**DOI:** 10.1101/2023.04.23.537997

**Authors:** Caleb A. Lareau, Yajie Yin, Jacob C. Gutierrez, Ryan S. Dhindsa, Anne-Sophie Gribling-Burrer, Yu-Hsin Hsieh, Lena Nitsch, Frank A. Buquicchio, Tsion Abay, Sebastian Zielinski, Robert R. Stickels, Jacob C. Ulirsch, Patrick Yan, Fangyi Wang, Zhuang Miao, Katalin Sandor, Bence Daniel, Vincent Liu, Quanli Wang, Fengyuan Hu, Katherine R. Smith, Sri V.V. Deevi, Patrick Maschmeyer, Slavé Petrovski, Redmond P. Smyth, William J. Greenleaf, Anshul Kundaje, Mathias Munschauer, Leif S. Ludwig, Ansuman T. Satpathy

## Abstract

Somatic variation contributes to biological heterogeneity by modulating cellular proclivity to differentiate, expand, adapt, or die. While large-scale sequencing efforts have revealed the foundational role of somatic variants to drive human tumor evolution, our understanding of the contribution of mutations to modulate cellular fitness in non-malignant contexts remains understudied. Here, we identify a mosaic synonymous variant (m.7076A>G) in the mitochondrial DNA (mtDNA) encoded cytochrome c-oxidase subunit 1 gene (*MT-CO1*, p.Gly391=), which was present at homoplasmy in 47% of immune cells from a healthy donor. Using single-cell multi-omics, we discover highly specific selection against the m.7076G mutant allele in the CD8^+^ effector memory T cell compartment *in vivo*, reminiscent of selection observed for pathogenic mtDNA alleles^1, 2^ and indicative of lineage-specific metabolic requirements. While the wildtype m.7076A allele is translated via Watson-Crick-Franklin base-pairing, the anticodon diversity of the mitochondrial transfer RNA pool is limited, requiring wobble-dependent translation of the m.7076G mutant allele. Notably, mitochondrial ribosome profiling revealed altered codon-anticodon affinity at the wobble position as evidenced by stalled translation of the synonymous m.7076G mutant allele encoding for glycine. Generalizing this observation, we provide a new ontogeny of the 8,482 synonymous variants in the human mitochondrial genome that enables interpretation of functional mtDNA variation. Specifically, via inter- and intra-species evolutionary analyses, population-level complex trait associations, and the occurrence of germline and somatic mtDNA mutations from large-scale sequencing studies, we demonstrate that synonymous variation impacting codon:anticodon affinity is actively evolving across the entire mitochondrial genome and has broad functional and phenotypic effects. In summary, our results introduce a new ontogeny for mitochondrial genetic variation and support a model where organismal principles can be discerned from somatic evolution via single-cell genomics.

## Main

The human body is comprised of thousands of distinct cell types with highly-specialized functions. Notably, the impact of pathogenic genetic variants on functional phenotypes may be pronounced in specific cell states but silent in others, indicative of specific demands of otherwise ubiquitous molecular processes, including translation and oxidative phosphorylation^3, 4^. Recent single-cell sequencing efforts have charted the effects of state-specific variant functions across human cell types, predominantly through studying common variant effects via single-cell expression quantitative trait loci mapping^5, 6^. Other approaches utilize single-cell genotyping to discern the transcriptional and epigenetic impact of somatic variants across different lineages in the hematopoietic system^7, 8^. While these and other studies enable a new annotation of functional variation in the human genome, our understanding of cell type-specific functional DNA variation remains underexplored.

Recently, we have introduced single-cell multi-omic methods detecting genetic variants in mitochondrial DNA (mtDNA) with concomitant measures of cell state, including accessible chromatin^9^, gene expression^10^, and/or protein abundances^11, 12^. While these methods have broadly enabled lineage tracing across human tissues, their application to congenital mitochondrial disorders has revealed previously unappreciated state-specific impacts of single nucleotide variation^1^ and large-scale mtDNA deletions (SLSMD) across human immune cells^2^. Noting mtDNA is present at approximately 100 copies per cell in lymphocytes, we demonstrated that distinct immune cell lineages differentially tolerate loss-of-function (LOF) mutations, including the notable purifying selection of the m.3243A>G mutation in T cells^1^. More recently, we have refined these observations and implicate CD8^+^ effector-memory T cells (CD8^+^ TEM) to be particularly vulnerable to LOF heteroplasmy^2^, suggesting that activation and expansion of naive CD8^+^ T cells require intact mitochondrial genetic integrity.

Here, we report the identification of a mosaic synonymous mutation, m.7076A>G (*MT-CO1*:p:Gly391=), in peripheral blood mononuclear cells (PBMCs) from a healthy donor. While we observe similar allelic heteroplasmy of this variant in all hematopoietic lineages, we observed a depletion of the m.7076G allele specifically in the CD8^+^ TEM compartment, reminiscent of the selection of diverse pathogenic mtDNA variants^1, 2^. We demonstrate that due to the limited diversity of the transfer RNA (tRNA) pool in mitochondria, this synonymous mutation requires translation via the super wobble effect, where a 5’ uracil in the anticodon can decode all four nucleotides in the 3’ codon. While capable of translation, the U-G wobble base pairing stalls mitochondrial ribosomes, thereby impeding CD8^+^ T differentiation. The principles of wobble-mediated translation elucidated from this mosaic variant enable a new ontogeny of synonymous mtDNA that broadly impact human genotypes and phenotypes. Specifically, we provide genetic evidence that germline and somatic synonymous mitochondrial variants undergo “codon optimization” in tumors, healthy tissues, and the germline across six different population-based cohorts (TCGA, IMPACT, PCAWG, GTEx, gnomAD, HelixMTdb). Comprehensive inter- and intra-species evolutionary analyses of the mitochondrial genome suggest that nucleotides in the wobble position experience accelerated evolutionary pressure between species, and human haplogroup-defining synonymous variation are enriched for variation consistent with preference for codon optimization. Based on population-level genotype associations from the UK BioBank (UKBB), synonymous variants that variably require wobble-dependent translation impact a variety of human traits and complex diseases. Altogether, we delineate a new ontology of functional synonymous variation altering codon syntax in the mitochondrial genome with broad relevance to human phenotypes, disease, and evolution.

## Results

### A mosaic synonymous mtDNA variant that is selected against in the CD8^+^ T cell compartment

Over the course of longitudinal profiling of the clonal dynamics of a healthy donor, we profiled PBMCs with the mitochondrial scATAC-seq assay (mtscATAC-seq^9, 13^) to simultaneously yield cell state and mtDNA genotyping information from the same single cells (**Fig. 1a**). Consistent with our prior observations from native hematopoiesis in healthy individuals^9^, we identified 183 somatic mtDNA variants with a median allele frequency of 0.04% in pseudobulk (**Fig. 1b; Methods**). Notably, we observed one variant, m.7076A>G, present at 47.3% heteroplasmy, suggesting this donor to be a chimera for this particular allele that likely arose during development. The m.7076A>G variant is a synonymous variant (p.Gly391=) in the mitochondrial cytochrome c oxidase subunit 1 (*MT-CO1*), a gene required for complex IV activity during oxidative phosphorylation (**Fig. 1c-d**). The majority of the 33,754 profiled cells had either 0% or 100% heteroplasmy (**Fig. 1c, Extended Data Fig. 1a**), a pattern that we previously observed for a non-coding mutation m.16260C>T in an independent healthy donor^11^. While m.7076A>G is present as a homoplasmic variant in 0.2%-0.9% of the population^14, 15^, the variant does not define a specific mitochondrial haplogroup. Still, approximately 1 in 15,000 individuals carries a highly heteroplasmic m.7076A>G allele (10-90% allele frequency) similar to this donor^14, 15^ (**Methods**).

**Figure 1.**
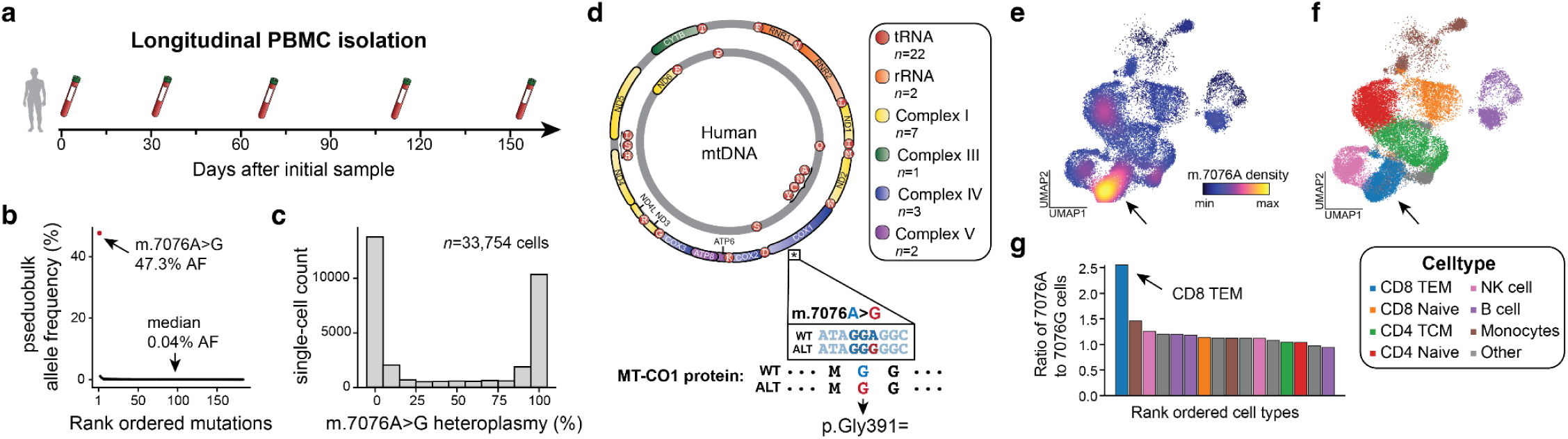
Identification of a mosaic synonymous mtDNA variant with a CD8^+^ T-cell restricted selection bias. **(a)** Schematic of longitudinal peripheral blood samples obtained from a healthy donor. A total of 5 draws spanning 150 days were taken and processed with the mtscATAC-seq assay. **(b)** Summary of somatic mtDNA mutations called from aggregated draws. The overall median heteroplasmy is noted, as well as the m.7076A>G allele that is present at a 47.3% pseudobulk heteroplasmy. **(c)** Distribution of single-cell heteroplasmy across all cells profiled for the m.7076A>G allele. **(d)** Schematic of the mitochondrial genome with genes contributing to indicated complexes of the respiratory chain (I, III, IV, and V) being color-coded. The asterisk under the *MT-CO1* genes denotes the position of the m.7076 allele. Annotation of the mutation, including the protein consequence (p.Gly391=) of the synonymous variant is shown below. **(e)** Uniform manifold approximation and projection (UMAP) of accessible chromatin profiles of PBMCs assayed via mtscATAC-seq colored by the density of the m.7076A (wildtype) allele. **(f)** Cluster annotation and cell type labeling of the same cells as in (e). The arrow indicates the CD8^+^ T effector memory (CD8^+^ TEM) population. **(g)** Ratio of homoplasmic cells with wildtype m.7076A to mutant m.7076G variants across indicated cell state clusters. The arrow highlights the CD8^+^ TEM cell state as the population with the greatest skew (p < 2.2e-16; binomial test).

Leveraging the chromatin accessibility modality of the mtscATAC-seq dataset, we utilized a dictionary learning strategy to infer the cell states for each cell that was concomitantly genotyped (**Methods**). Unlike in any other cell state, we observed a marked depletion of cells homoplasmic for the mutant m.7076G allele in a subset of CD8^+^ TEM (**Fig. 1e,f, Extended Data Fig. 1b-d**). In other words, CD8^+^ TEMs had a marked increase of cells with the wildtype m.7076A allele, suggestive of lineage-specific selection pressure against cells with the mutant m.7076G allele (**Fig. 1g**). Notably, these findings mirror our recent observations of purifying selection against pathogenic mtDNA mutations in CD8^+^ TEM cells in individuals with congenital mitochondriopathies, including patients with Mitochondrial Encephalopathy, Lactic Acidosis, and Stroke-like episode (MELAS)^1^ and Pearson Syndrome^2^, which is driven by the distinct demand for OXPHOS capacity during T cell proliferation and differentiation after activation^16, 17^.

### Altered T-cell state-specific gene expression due to m.7076A>G

As synonymous variants alter the sequence of DNA but retain the amino acid sequence of the encoded protein, these alleles do not tend to have a functional impact and are typically annotated as ‘silent’ mutations. However, differential effects of distinct tRNAs decoding synonymous codons and alterations in enzyme structure and function due to synonymous mutations have been reported^18^, including in human tumor progression^19, 20^. Thus, we hypothesized that the synonymous variant m.7076A>G could incur a loss of function phenotype resulting in selective pressure during T cell differentiation analogous to known pathogenic mtDNA variants. To examine the possibility of this mutation impacting *MT-CO1* mRNA expression, we performed single-cell RNA-seq (scRNA-seq) profiling of 17,337 T-cells derived from PBMCs derived from this donor. We verified the specific depletion of the m.7076G allele in the CD8^+^ TEM compartment, but not other T cell subsets (**Fig. 2a**). As a recent study attributed functional synonymous variation in yeast to differences in gene expression^21^, we compared the abundance of the *MT-CO1* transcript between cells with homoplasmy for either the wildtype or mutant allele (**Methods**). Within any of these cell subsets, we did not observe significant differences in expression, suggesting that the synonymous variant does not impact *MT-CO1* stability or expression (**Fig. 2b**).

**Figure 2.**
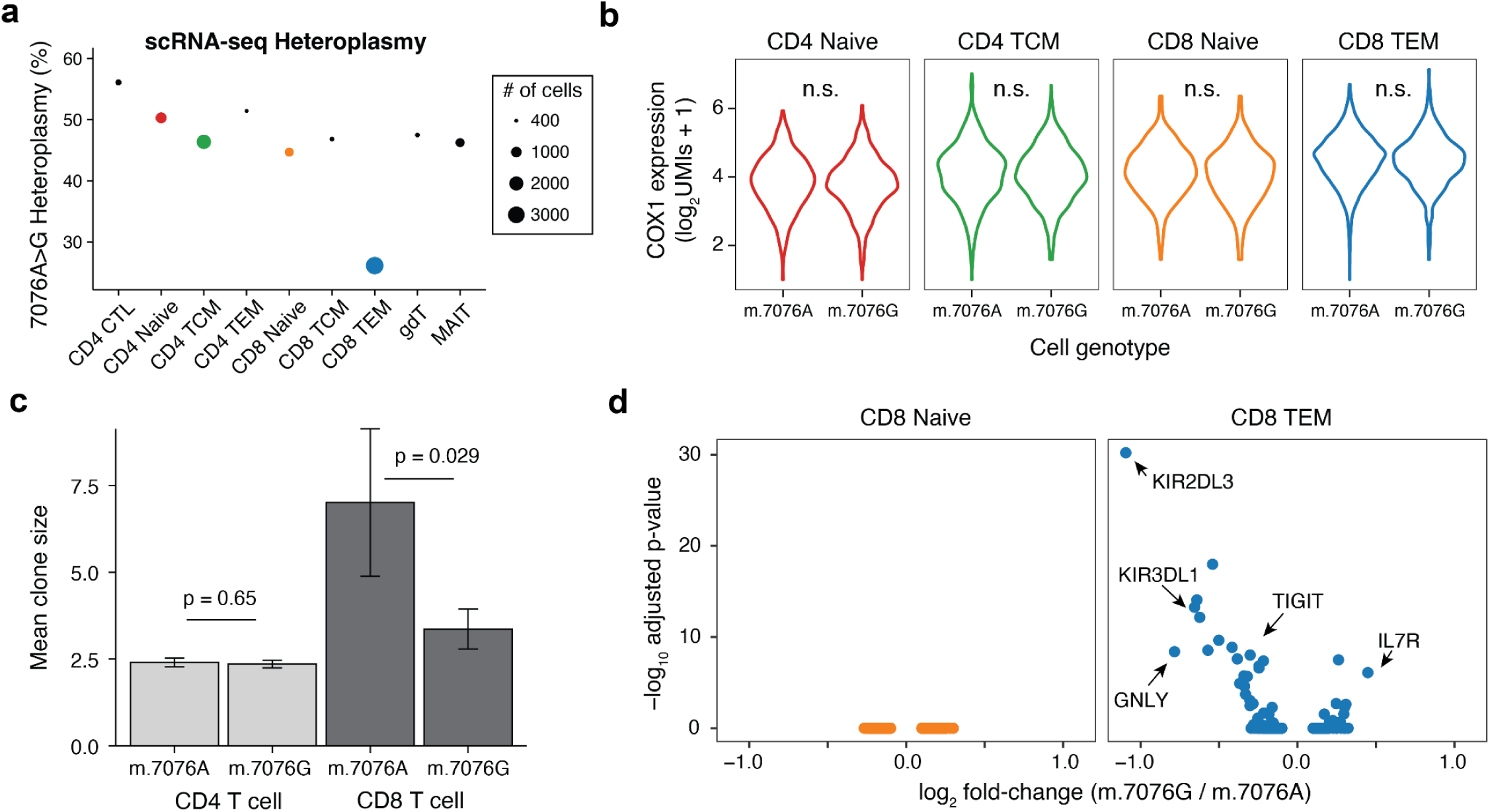
Stable expression of *MT-CO1* transcript but altered clone size of CD8^+^ effector memory T cells carrying the mutant m.7076G allele. **(a)** Heteroplasmy of m.7076A>G in indicated T cell subpopulations based on scRNA-seq. The size of each dot is scaled by the abundance of cells in each cell state. **(b)** Comparison of *MT-CO1* expression across indicated cell states stratified by the m.7076A and G alleles. P-values from a Wilcox test comparing log_2_ *MT-CO1* UMI counts per cell were not significant at type I error of 0.05. **(c)** Comparison of TCR clone sizes (number of cells per clone) between CD4^+^ and CD8^+^ T cells with homoplasmic m.7076A or m.7076G alleles. P-values are shown for a Wilcox test comparing clone sizes which were significant at type I error of 0.05. **(d)** Differential gene expression of all genes between cells with homoplasmic wildtype m.7076A vs. mutant m.7076G within the indicated CD8^+^ T cell compartments. 0 genes were differentially expressed in naive CD8^+^ T cells whereas 32 were differentially expressed in CD8^+^ TEM cells, including the 5 highlighted in the text. No other differentially expressed genes were observed in other T cell subsets

Utilizing concomitant single-cell T-cell receptor (TCR) sequencing, we examined clone sizes of CD4^+^ and CD8^+^ T cells with either m.7076 allele. As expected, TCR clones were effectively mutually exclusive for either m.7076 allele, confirming an early developmental origin of the mutant allele prior to TCR diversification (**Extended Data Fig. 2a**). We observed a diminished clone size in the CD8^+^ but not CD4^+^ T cell compartment in cells with the mutant m.7076G allele (**Fig. 2c**), suggesting reduced fitness or proliferative capacity of mutant CD8^+^ T cells to clonally expand. Further, we performed differential gene expression analyses across T cell subsets between wildtype and mutant m.7076 cells. Whereas all CD4^+^ subsets and CD8^+^ naive subsets had no differentially expressed genes between cells with distinct genotypes, we observed 32 differentially expressed genes between wildtype and mutant CD8^+^ TEM cells (**Fig. 2d**). While cytotoxic genes including *GNLY*, *KIR2DL3*, and *KIR3DL1* were down-regulated, *IL7R*, a marker of naive T cells, was more highly expressed in cells with the mutant m.7076G allele. Additional unsupervised analyses revealed distinct cell states within the CD8^+^ TEM compartment, including highly clonal wildtype m.7076 allele populations expressing specific TCRs (**Extended Data Fig. 2b-c**). However, these highly expanded clones can not alone explain the observed depletion of the mutant allele in CD8^+^ TEM (**Extended Data Fig. 2d**; **Methods**). Though the mutant m.7076A>G allele does not preclude the possibility of CD8^+^ TEM differentiation, we conclude that mutant cells are at a competitive disadvantage to expand and acquire fully differentiated cytotoxic T cell-like phenotypes as evidenced by functional gene expression defects.

### Refinement of selected T cell phenotypes

To further investigate the CD8^+^ T cell deficiencies incurred by this variant, we stimulated PBMC-derived T cells and examined cell phenotypes using flow cytometry and ATAC with Selected Antigen Profiling via sequencing (ASAP-seq) that extends the mtscATAC-seq to co-quantify surface marker profiles via oligo-barcoded antibodies (**Extended Data Fig. 3a**). Leveraging the multi-modal measurements, we again observed a focalization of the wildtype m.7076A allele in a specific CD8^+^ T cell cluster (**Extended Data Fig. 3b,c**). Interestingly, this cluster is characterized by high KLRG1 surface protein expression but depleted of IL7R protein, consistent with the transcriptomics data (**Extended Data Fig. 3d,e**). Gene scores showed highly accessible chromatin in the *ZEB2*, *IFN-γ*, and *KLRD1* loci, suggestive of a CD8^+^ short-lived effector cell (SLEC)-like population that is 1.3x enriched for cells with the wildtype m.7076A allele compared to other T cell subsets (**Extended Data Fig. 3f,g;** *p*=1.6×10^-11^; binomial test). Thus, our T-cell culture experiments confirm mutant m.7076A>G cells to be impaired to attain specific CD8^+^ T cell states *in vitro*.

**Figure 3.**
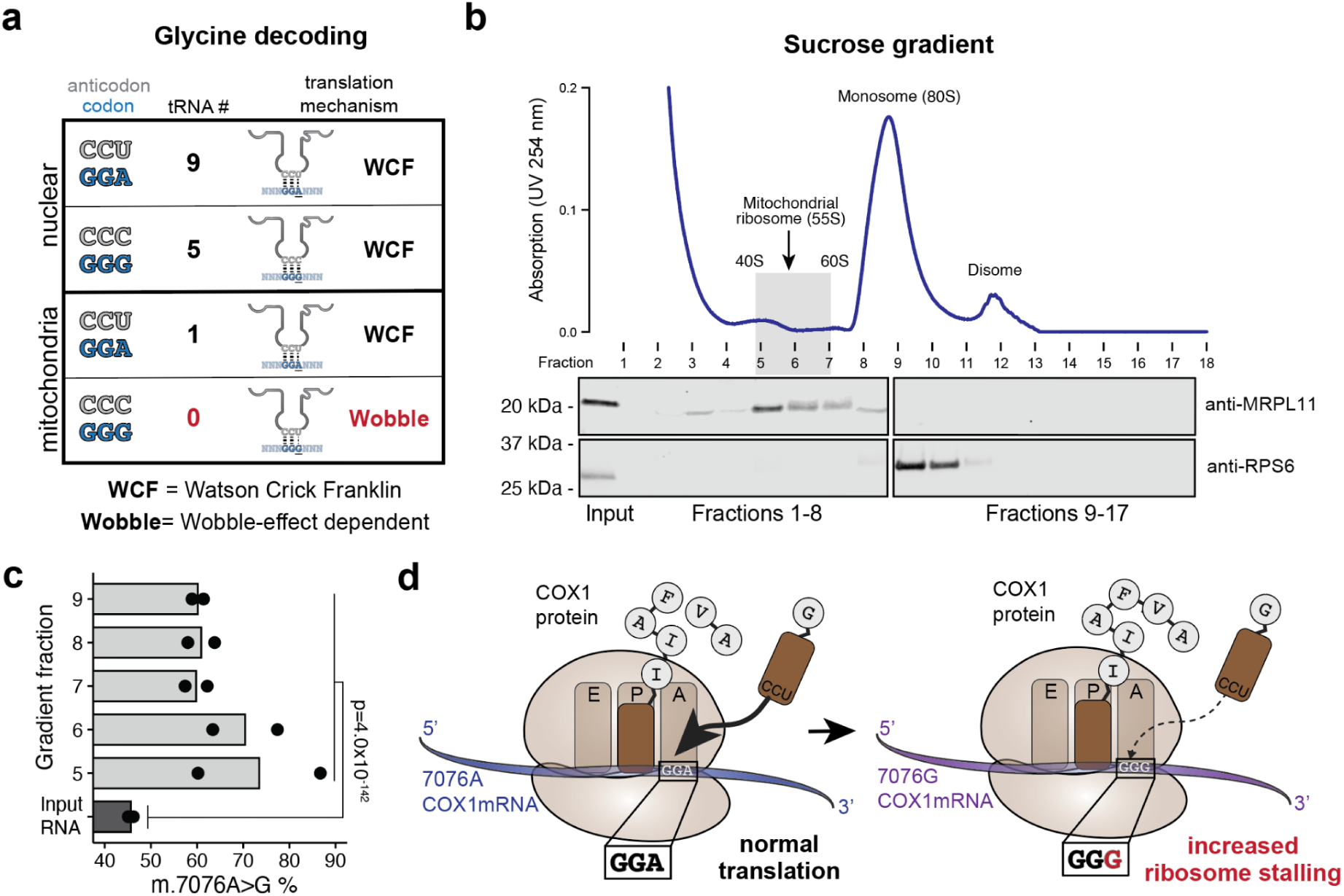
Mitochondrial ribosome profiling reveals translational stalling of the mutant m.7076A>G allele. **(a)** Summary overview of codon, anticodon, tRNA, and codon:anticodon recognition mechanisms for glycine in the human nuclear and mitochondrial genomes. **(b)** Polysome profile following sucrose gradient and western blots of isolated fractions for mitochondrial ribosome profiling. MRLP11 and RPS6 were blotted to identify enrichment of mitochondrial and cytoplasmic ribosomes, respectively **(c)** Summary of heteroplasmy from ribosome profiling libraries (fractions 5-9, see panel (b)) showing a relative increase of the mutant m.7076A>G allele in ribosomal bound fractions versus input RNA. Statistical significance was determined using a Fisher’s exact test of 7076A and G read counts summed between replicates. **(d)** Schematic of the functional effect of the synonymous m.7076A>G variant. Due to decreased codon:anticodon affinity of the m.7076A>G allele, there is an increase in stalling of the *MT-CO1* transcript, prohibiting effective translation.

### Altered wobble-dependent translation of m.7076A>G

Given no alterations in *MT-CO1* transcript abundance, we hypothesized a post-transcriptional mechanism to be responsible for the mutant functional effects. As synonymous codons may be decoded by independent tRNAs, we revisited the distinct pool of tRNAs in mitochondrial translation compared to nuclear-encoded tRNAs for cytoplasmic protein biosynthesis. Specifically, the nuclear genome redundantly encodes multiple tRNA loci for all possible codons, but only 22 tRNAs are ordinarily transcribed in the human mitochondrial genome. Thus, only a single glycine-specific tRNA (tRNA^Gly^) is encoded in the mitochondrial genome to decode p.Gly391 via the wildtype GGA codon (m.7076A) or the mutant GGG codon (m.7076A>G; **Fig. 3a**). As such, decoding via the single mt-tRNA^Gly^ of the wildtype GGA allele occurs via canonical Watson-Crick-Franklin (WCF) base-pairing but requires wobble-dependent translation of the mutant GGG codon (**Fig. 3a**). Although the GGG codon is relatively depleted in the mitochondrial genome compared to the nuclear genome (**Extended Data Fig. 4a,b**), this codon occurs at 34 positions within the 13 peptide-encoding open reading frames, suggesting the mitochondrial translational machinery to be generally capable of decoding this codon. Based on our cell state mapping experiments, we hypothesized that the substantial requirement for oxidative phosphorylation during CD8^+^ T cell activation, differentiation, and proliferation may present a distinct metabolically vulnerable state toward CD8^+^ TEM cell state transition that is sensitive to even subtle changes in oxidative phosphorylation capacity.

**Figure 4.**
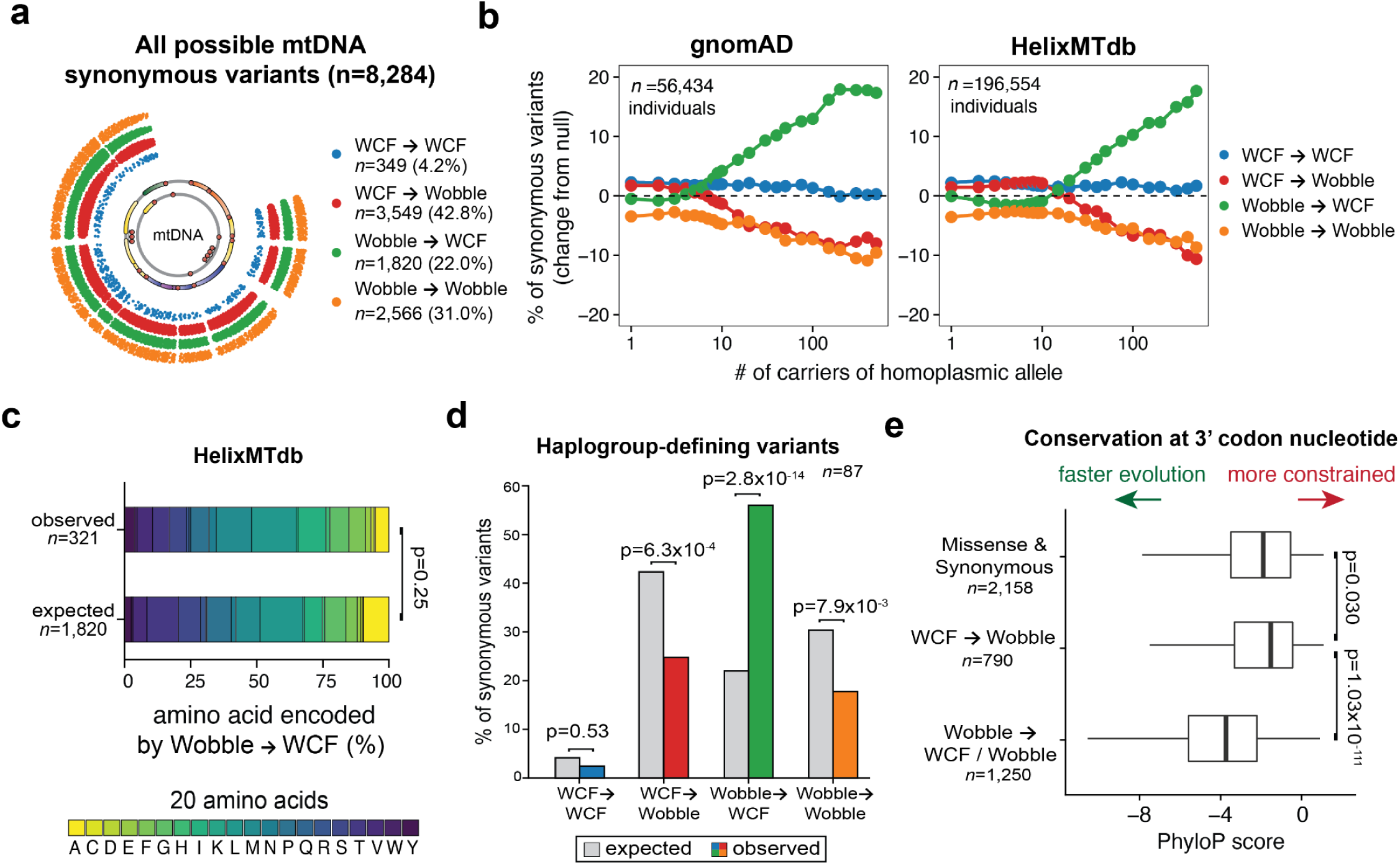
A new ontogeny of synonymous mtDNA mutations reveals patterns of human germline variation mediated by wobble-dependent translation. **(a)** Classification of all 8,284 possible synonymous mtDNA variants based on Watson-Crick-Franklin (WCF) or wobble-dependent base-pairing at either the reference or alternative allele. **(b)** The abundance of each synonymous variant class at different thresholds for the number of carriers for two population-scale sequencing databases. The difference in the percent of all synonymous variants is plotted for the four variant classes (null: dotted line at 0). **(c)** The proportion of the amino acid encoded by the codons relevant for the Wobble→WCF variants. The observed variants represent Wobble→WCF present in at least 100 individuals from HelixMTdb. The test statistic is a Chi-squared test for the number of observed Wobble→WCF variants encoding each amino acid. **(d)** Comparison of % observed (color) versus expected (grey) for 87 haplogroup-defining, synonymous mtDNA mutations. P-values represent the statistical significance of a two-sided binomial test statistic. **(e)** Inter-species conservation at wobble-position nucleotides in mitochondrial codons. Reference alleles that can be mutated to different outcomes are specified and grouped with the number of wobble-position codons in each class noted. P-values represent a Wilcoxon test.

To examine this possibility, we stimulated PBMCs in the presence of αCD3/αCD28 beads and IL-2 and assessed mitochondrial translational efficiency (**Methods**). We isolated fractions along a sucrose gradient to enrich mitochondrial ribosomes and prepared libraries for mitochondrial ribosome profiling by sequencing (MitoRiboSeq; **Fig. 3b**: **Methods**). Hypothesizing the altered codon syntax of the mutant m.7076A>G allele to impair translational decoding, the mutant G allele would be relatively enriched in the ribosome-protected fragments compared to the total mRNA input. Indeed, across all 10 fractions (5 per biological replicate), we computed a translation pause ratio, defined as the fraction of reads from ribosome profiling over the RNA-seq libraries, and observed that the mutant m.7076A>G allele was enriched over the input material (**Fig. 3c, Extended Data Fig. 4c**, mean 33% increase; p=4.0×10^-142^), which was further replicated in an independent experiment (**Extended Data Fig. 4d**, 3 additional replicates; mean 37% increase; p=3.1×10^-24^). Specifically, the wildtype and mutant alleles were present at similar transcript levels (54.3% m.7076A vs. 45.7% m.7076G; ratio 1.19) compared to the mtscATAC-seq based genotyping results (47.3% m.7076G) further confirming *MT-CO1* mRNAs abundance to be relatively unaltered. However, among ribosomal protected fragments, the mutant allele was substantially more abundant (39.1% wildtype m.7076A vs. 60.9% mutant m.7076A>G; ratio 0.64), strongly evident of translational stalling around the mutant codon (**Extended Data Fig. 4c**). Thus, the limited diversity of the tRNA pool prohibits efficient translation of the mutant m.7076A>G allele, otherwise requiring the super-wobble effect to effectively decode the mutant GGG codon. As a consequence, translation of *MT-CO1* is stalled via a mechanism consistent with reports of super-wobble model systems^22^, thereby resulting in an effective partial LOF that becomes particularly restricting during the metabolically demanding process of CD8^+^ T cell differentiation.

### A new ontogeny of synonymous mtDNA variation

Given the overall restricted mitochondrial tRNA pool, we evaluated the potential impact of synonymous mtDNA variation beyond the mutant m.7076A>G allele and considered all possible synonymous mutations across the mitochondrial genome, totaling 8,284 variants across 13 polypeptide-encoding genes (**Fig. 4a**). In our ontogeny, we annotated variants based on whether the reference and alternate alleles utilize canonical WCF or wobble base-pairing, leading to four possible classifications for synonymous variants: WCF→WCF, WCF→Wobble, Wobble→WCF and Wobble→Wobble (**Fig. 4a; Methods**). From our annotation, we observed 48% of codons in the mitochondrial genome require wobble-dependent translation; however, to our knowledge, potential functional effects of codon utilization in the human mitochondrial genome have not been reported. To examine this annotation, we first examined public MitoRiboSeq data^23^ from two cell lines (HEK293 and HeLa) and compared homoplasmic variants specific to each line (**Methods**). For two additional mtDNA variants, we observed sufficient coverage to examine differences in MitoRiboSeq coverage and confirmed evidence of increased stalling due to a wobble-dependent base pairing but not a WCF pairing (**Extended Data Fig. 4e,f**).

To assess the impact of this mitochondrial genome-wide annotation, we queried gnomAD^24^ and HelixMTdb^14^, two comprehensive databases of germline variation in humans, where mtDNA variation has been called for 56,434 and 196,554 healthy individuals, respectively. While the simple occurrence of mtDNA variants (observed in ≥1 individual) mostly followed the expected proportions, the more common synonymous variants (present in multiple individuals) were enriched for variants that increased codon affinity (i.e., Wobble→WCF; “more optimal base-pairing”) and depleted for variants that reduced codon affinity (i.e., WCF→Wobble or Wobble→Wobble; “less optimal base-pairing”; **Fig 4b; Extended Data Fig. 5a,b**). This pattern of codon-optimizing variation increasing in the population was replicated between both gnomAD (p<2.2×10^-16^; Mann-Kendall Trend Test) and HelixMTdb (p=2.38×10^-7^; Mann-Kendall Trend Test). Wobble→WCF variants in at least 100 donors from HelixMTdb were observed in codons for all amino acids at proportions that did not deviate from the expected proportions (**Fig 4c;** p=0.25; Chi-squared test), indicating that this selection for codon optimality impacted all amino acids. In total, our analyses demonstrate that variants impacting wobble-dependent translation are not constrained, consistent with recent work^25^, but accumulate in a way that optimizes codon:anticodon affinity, indicative of positive selection at the population level.

**Figure 5.**
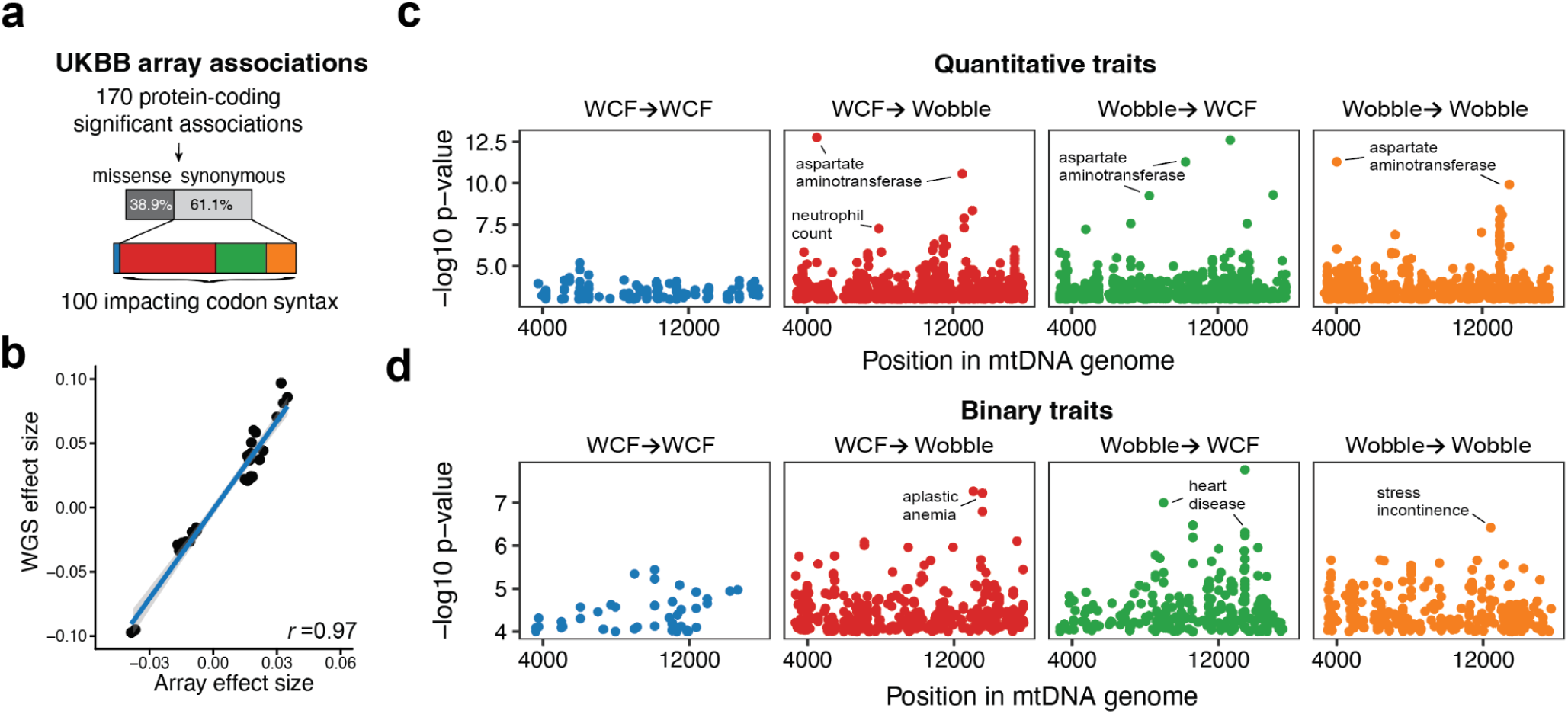
Complex human phenotypes are associated with synonymous mtDNA mutations. **(a)** Reanalysis of statistically significant associations in protein-coding genes between mtDNA variants measured by a genotyping array and complex traits assayed in the UKBB^29^. **(b)** Correlation of effect sizes for common variants estimated from WGS (y-axis; computed in this study) and prior array-based work^29^. Only variant-quantitative trait pairs that were statistically significant in the array-based study^29^ are noted, and the Pearson correlation between the effect sizes is shown. **(c)** Manhattan plots of all synonymous mtDNA variants stratified by wobble-dependent ontogeny using a large biobank of whole-genome sequencing data. **(d)** Same as in (c) but for binary traits. Each dot represents a unique trait-genotype association with a p-value <1×10^-4^.

### mtDNA wobble codons experience evolutionary selection

Next, we investigated if mitochondrial genomes displayed evolutionary evidence of codon selection. To examine this, we considered both intra-species (**Fig. 4d**) and inter-species evolution (**Fig. 4e**). We identified 87 common synonymous variants linked to the phylogeny of human mitochondrial haplogroups^26, 27^ and examined their proportions in light of our new ontogeny (**Methods**). Here, we observed a >2.5 fold increase of Wobble→WCF variants (observed: 56.3%; expected: 22.0%; p=2.8×10^-14^), suggesting that variation linked to human mtDNA haplotypes for increased WCF pairing compared to the most recent common human ancestor is an ongoing evolutionary process. We then considered inter-species annotations of conservation, requiring a per-position analysis rather than a per-variant analysis (**Methods**). Considering all three possible variant outcomes for a hypothetically mutated position, we classified each of the wobble-position nucleotides in the mitochondrial genome into three categories: positions that could variably lead to missense or synonymous variants (*n*=2,158), WCF reference alleles that become wobble in all scenarios (*n*=790), or wobble reference alleles that when mutated are either wobble or WCF (*n*=1,250). Using both a 20 and 100 species phyloP score^28^, we assessed the evolutionary conservation of each codon class, confirming minimal constraint across any nucleotide consistent with prior results^25^ (**Fig. 4e**). Notably, we observed a significant decrease in phyloP score specifically for positions that require the wobble effect for translation in the reference genome, suggesting that loci requiring the wobble effect are under accelerated evolution compared to already optimal WCF pairings (**Fig. 4e**; p=1.03×10^-111^; Wilcoxon test). These results indicate that though human mtDNA possesses abundant codons that require wobble-dependent translation, the ongoing evolution of the human mtDNA genome is selecting for more optimal WCF-compatible base-pairing.

### Complex trait associations

We hypothesized that our classification of synonymous mtDNA variants would enable a new interpretation of results from genome-wide association studies. To assess this, we reanalyzed association summary statistics from a recent study that examined the landscape of complex phenotypes from 488,377 individuals array-genotyped for 265 mtDNA SNVs in the UK Biobank while adjusting for nuclear/haplotype associations^29^. Notably, most of the significantly associated SNPs in protein-coding genes were synonymous variants (104 of 170 or 61.1% of significant SNP-trait associations; **Fig. 5a**). 100 of the 104 synonymous variants altered codon syntax, more than a random sampling of variants (p=0.038; proportion test), including WCF→Wobble variants that confer risk for bipolar disorder (m.13117A>G; *MT-CYB;* FDR=0.012) and type 2 diabetes (m.8655C>T; *MT-ATP6*; FDR=0.046). While prior studies of these synonymous variants suggested they tag a separate causal variant^29^, we interpret these associations to reflect the altered codon syntax impairing the effective translation and/or structure/function^30^ of mitochondrial proteins, thereby impacting specific cell state function and differentiation, resulting in a heightened predisposition for a myriad of complex human traits.

As this mtDNA association analysis relied on variants present in a genotyping array, we leveraged the detection of the mitochondrial sequence information in whole-genome sequencing data to assess associations between complex traits and rare mtDNA variants (**Methods**). After verifying consistent effect estimates between the genotyping array and our WGS sequencing analyses (**Fig. 5b**), we performed systematic association studies between 2,786 commonly observed synonymous mtDNA variants with 1,656 quantitative phenotypes (**Fig. 5c**) and 18,688 binary phenotypes (**Fig. 5d**) from 171,673 UKBB participants (**Methods**). Using a permutation method to determine the statistical significance of our associations (**Methods**), we identified 492 variant-trait associations from the synonymous variants, only 12 of which were WCF→WCF. Further, WCF→WCF variants had a distinct distribution of test statistics compared to the other classes of synonymous variation altering codon affinity (p=9.6×10^-7^; Wilcoxon test), suggesting germline variants that alter mtDNA codon syntax may be broadly associated with complex human phenotypes.

We observed the most significant associations between several rare variants altering mitochondrial codon affinity and peripheral blood cell traits, including neutrophil count with m.7235C>A (Wobble→WCF; *MT-CO1*; p=2.7×10^-8^) as well as circulating metabolites and enzymes, including aspartate aminotransferase levels with m.4529A>T (WCF→Wobble; *MT-ND2*; p=1.7×10^-13^), m.4023T>C (Wobble→WCF; *MT-ND5;* p=5.1×10^-12^), and m.10238T>C (Wobble→Wobble; *MT-ND1*; p=5.2×10^-12^; **Fig. 5c**), which may be linked to mitochondrial metabolic activity in the liver^31^. Among disease phenotypes, we observed significant associations with multiple genitourinary diseases (m.10598A>G; p=3.3×10^-7^; *MT-ND4L;* Wobble→WCF) and aplastic anemia (m.13557A>G; p=5.9×10^-8^ *MT-ND5;* WCF→Wobble; **Fig. 5d**). For these highlighted variants, we hypothesize that a functional effect in the mitochondria may be realized in a restricted cell state, analogous to the CD8^+^ T cell population implicated in the m.7076A>G mutation. In sum, our array- and sequencing-based analyses corroborate that our new annotation of wobble-dependent mtDNA variation broadly impacts complex traits via modulating genes encoded in the mitochondrial genome.

### Codon syntax is somatically optimized in human tumors and tissues

Systematic analyses of tumor sequencing data have suggested that translational control of the nuclear genome via codon usage can substantially modulate oncogenic processes^20, 32^. Thus, we sought to examine a potential link between mtDNA synonymous mutation classes and somatic occurrence in tumors, we analyzed a total of 2,197 synonymous mtDNA mutations across three sequencing cohorts (**Fig. 6a**). Notably, we observed a significant depletion of WCF→Wobble variants (“less optimal”; observed: 30.3%; expected: 42.8%; p=2.3×10^-32^; proportion test) and a corresponding increase of Wobble→WCF (“more optimal”; observed: 35.9%; expected: 22.0%; p=1.4×10^-55^; proportion test) across all cohorts. We stratified all possible synonymous mutations into four major classes based on the chemical structure of the wobble base to its paired tRNA (**Fig. 6b; Methods**). Many mitochondrial tRNAs are post-transcriptionally modified at the wobble position in the anticodon, leading to non-canonical nucleotides like queuosine, or uracil (super-wobble)^33^. Irrespective of tRNA class, we again observed a consistent depletion of WCF→Wobble variants in the pan-cancer datasets (**Fig. 6b**), indicating this ontogeny of functional synonymous mtDNA likely impacts all amino acids and tRNA classes roughly equivalently.

**Figure 6.**
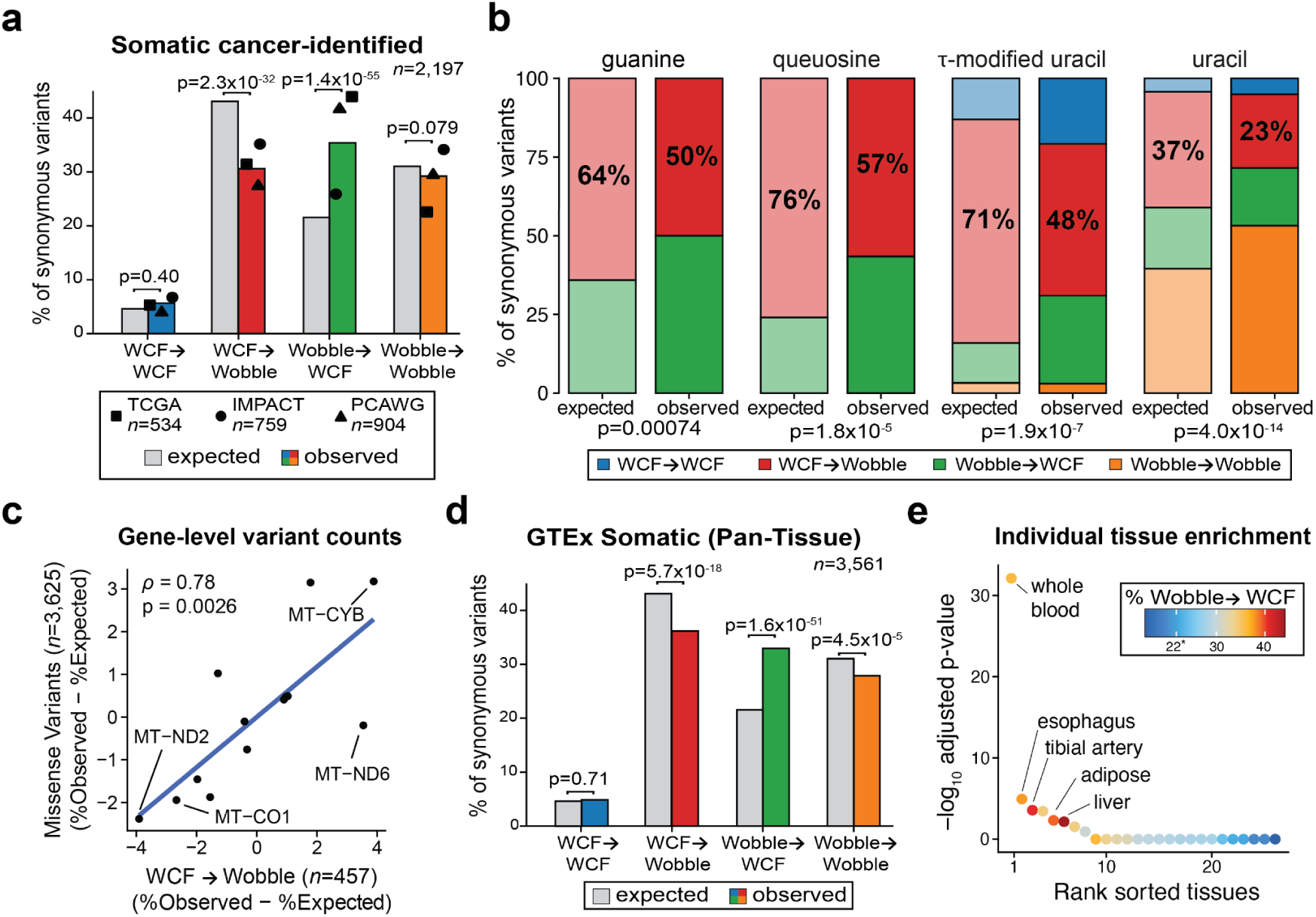
Wobble-dependent translation reveals patterns of synonymous somatic mtDNA variation in tumors and healthy tissues. **(a)** Comparison of observed (color) versus expected (grey) somatic synonymous mtDNA mutations based on three tumor atlas sequencing cohorts (TCGA, IMPACT, PCAWG). P-values represent the statistical significance of a two-sided binomial test statistic. **(b)** Stratification of synonymous variants by tRNA base in the anticodon wobble position. Faded colors represent the expected numbers of mutations under a null model. Percents represent the fraction of mutations that are WCF→Wobble. Chi-squared test of association between observed and expected mutation values based on numbers of potential or observed mutations in each synonymous mutation class are shown below each pair of bar plots. **(b)** Gene-level abundances of missense and WCF→Wobble variants comparing expected (based on all possible variants) versus observed variants across the entire dataset. The Pearson correlation and p-value from the Pearson correlation test are noted. **(d)** Same as in (a) but for somatic mutations identified in the mtDNA genome in one or more non-cancerous GTEx tissues^13^. **(e)** The proportion of synonymous variants that are Wobble→WCF within individual tissues. 22% (noted by the asterisk) is the abundance under the null. Tissues are rank-ordered by statistical significance using the p-value from a two-sided binomial test.

As mutations in specific mitochondrial genes/complexes may modulate clinical outcome^34^, we examined the abundance of WCF→Wobble variants (“less optimal”) and missense variants at a gene level compared to a null model (**Methods**). We hypothesized that the per-gene mutational burden of these classes of variants might be correlated as they may result in partial loss of function phenotypes either due to protein-level changes or decreased translational efficiency. After quantifying 3,625 unique missense variants and 457 WCF→Wobble variants across our compendium of somatic tumor mtDNA mutations, we observed a strong concordance between these two classes of genetic variants; for example, *MT-CYB* was frequently mutated while *MT-CO1* and *MT-ND2* mutations were consistently depleted (**Fig. 6c**). This disparity is consistent with their differential roles in mitochondrial biology. Specifically, mild disruption of complex III (including *MT-CYB*) can be beneficial for tumor growth whereas mutations in genes in complex IV (e.g., *MT-CO1*) and complex I (e.g., *MT-ND2*) were unfavorable^34, 35^. These results reflect a broad model of somatic “codon optimization” in tumors, consistent with reports of enhanced OXPHOS activity to be favorable during oncogenesis^34^, but noting that optimal translation rates can variably impact protein function^18^.

As this somatic codon optimization may be occurring in clonally expanded healthy tissues in addition to tumors, we annotated cross-tissue somatic mtDNA variation that we previously identified from the GTEx consortium^13^. Here, we observed a global enrichment of Wobble→WCF variation across all tissues (p=1.6×10^-51^; proportion test; **Fig. 6d**). Examining individual tissues, we observed enrichments of codon-optimizing variation in seven tissues (adjusted p<0.05; proportion test), including whole blood, esophageal, tibial artery, adipose, and liver tissues (**Fig. 6e**), noting that this analysis is impacted by the overall prevalence of mutations (of which whole blood was most abundant). We replicated the enrichment of somatic Wobble→WCF variants in peripheral blood of somatic mtDNA variation with single-cell sequencing (**Extended Data Fig. 6a**)^9^. We conclude that mutations in mitochondrial genomes experience selective pressure during tissue and tumor evolution, particularly in metabolically demanding cell states or cell state transitions, broadly leading to mitochondrial codon optimization.

### Murine mtDNA corroborates functional effect

As eukaryotic mtDNA were derived from a common ancestor, we hypothesized that our novel synonymous variant ontogeny should generalize to other species, including mice. We similarly annotated each of the 7,957 possible synonymous variants in the mouse GRCm39 reference genome in light of wobble-mediated transition (**Extended Data Fig. 7a**). This annotation broadly corroborated our inferences in humans, including the wobble nucleotides experiencing faster evolution than variants decreasing codon affinity or leading to missense mutations (**Extended Data Fig. 7b; Methods**). Next, we examined the somatic accumulation of synonymous mtDNA variation from pol-γ mutant mice^36^ and in healthy aged mice^37^ (**Extended Data Fig. 7c,d**). Comparison of these mice revealed a notable disparity where pol-γ mutant mice had enrichment of WCF→Wobble variants (‘less optimal’; observed: 57.1% expected: 52.2%; p=0.00094; proportion test) whereas healthy aged mice had >2x enrichment of Wobble→WCF variants (‘more optimal’; observed: 43.8%; expected: 20.9%; p≈0; proportion test), a result consistent with the somatic occurrence in healthy human tissues via GTEx analyses (**Fig. 6d**). As the pol-γ mutant mice have deficient proof-reading capacities in the polymerase required for mitochondrial DNA replication, the increased accumulation of somatic mutations in these mice have been tied to distinct organismal phenotypes, including erythroid dysplasia and impaired lymphoesis^38^. Thus, we suggest that diminished codon affinity via synonymous variation may contribute to the functional effects of pol-γ mutant mice. These results culminate in a consistent model where mitochondrial genome syntax is frequently somatically optimized during tumorgenesis, germline variation, and clonal expansion in healthy tissues across multiple species.

## Discussion

Mapping somatic DNA mutations across healthy tissues revealed hundreds of somatic mutations that undergo positive selection in variable physiologic contexts^39–41^. However, these mutations are infrequently shared between multiple tissues, suggesting that many somatic variants contribute to cell state heterogeneity in a context-specific manner^42^. Here, we characterized a chimeric somatic mtDNA variant m.7076A>G in the peripheral blood of a healthy individual to discern a functional role for a synonymous mutation. Consistent with our prior work of purifying selection against pathogenic mtDNA variants^1, 2^, cells harboring the m.7076G allele are relatively less fit than their wild-type m.7076A counterparts, which limits their clonal expansion, cytotoxic profile, and *in vivo* abundance. Ultimately, our characterization of the wobble-dependent functional effect of this allele enables a previously underappreciated annotation of the mitochondrial genome that was corroborated by comprehensive population genetics and evolutionary analyses.

Though reports in model organisms have described nuclear-encoded tRNA transfer to the mitochondria to compensate for deficiencies of the mitochondrial tRNA pool, to the best of our knowledge, no evidence of functional nuclear tRNAs in the mitochondria has been observed in humans. The pathogenicity of loss-of-function mtDNA-encoded tRNA mutations in MELAS (m.3243 in a leucine tRNA), MERRF (m.8344 in lysine tRNA), and other mitochondriopathies further suggest that lack of meaningful nuclear tRNA contribution to mitochondria in humans^43^. Thus, the translation of the 13 mtDNA-derived polypeptides utilizes only the 22 tRNAs encoded in the mitochondrial genome. Consistent with this concept, m.7076A>G requires wobble-dependent translation via mt-tRNA^Gly^ at the *MT-CO1*:p.Gly391 polypeptide, which occurs but leads to ribosome stalling and diminished CD8^+^ T cell fitness. Notably, this variant does not impact the abundance of the *MT-CO1* mRNA in any profiled cell state, providing an independent and distinct mechanism from a recent report of functional synonymous variation that impacts mRNA stability in yeast^21, 44–46^. Critically, using population-level somatic and germline sequencing data, we conclude that the functional effect mediated by wobble-dependent mitochondrial translation is not restricted to the m.7076A>G but may variably impact the 8,284 synonymous variants in the mitochondrial genome.

In bacteria, selection for codon:anticodon matching is strongest in species that are the fastest growing^47^. Similarly, our inferences of human cell states are associated with rapid proliferation, specifically CD8^+^ TEM, which is biased toward optimized codon syntax. Noting that mitochondria emerged from bacterial ancestors via endosymbiosis^48^, our data convey a model where similar selective pressures in mitochondria may emerge in states of high metabolic demands, including CD8^+^ T cell differentiation/activation or with even small competitive advantage contributing to tumor development and tissue evolution. Analysis of germline sequencing data suggests that while synonymous variation in the mitochondrial genome is not under constraint within individuals^24, 25^, we observe associations indicating that codons requiring the wobble effect for translation experience accelerated evolutionary selection and are enriched for haplotype-defining variation that results in codon-optimized mtDNA genomes. In other words, while wobble-dependent translation does not lead to notable LOF in the germline, cellular fitness at the organismal level appears to be under positive selection for codon-optimizing variation. Thus, our inferences from examining the association of a somatic mtDNA variant within an individual generalize to species-level evolutionary principles.

Taken together, our results demonstrate that selection of mtDNA occurs within healthy individuals, emphasizing the potential of non-germline mtDNA variation impacting specific cell states. The extension of our inferences of functional effects of synonymous variation within the mitochondrial genome to haplogroup-defining variants and large population-based cohorts suggests that altered codon syntax plays a broad but previously unappreciated role in modulating traits, disease phenotypes, and human mitochondrial evolution. With the continued application of single-cell DNA sequencing with concomitant cell state characterization, we anticipate that our multi-omics approach will continue to reveal unique insights into the contribution of DNA variation to human phenotypes in distinct cellular and evolutionary contexts.

**Extended Data Fig. 1.**
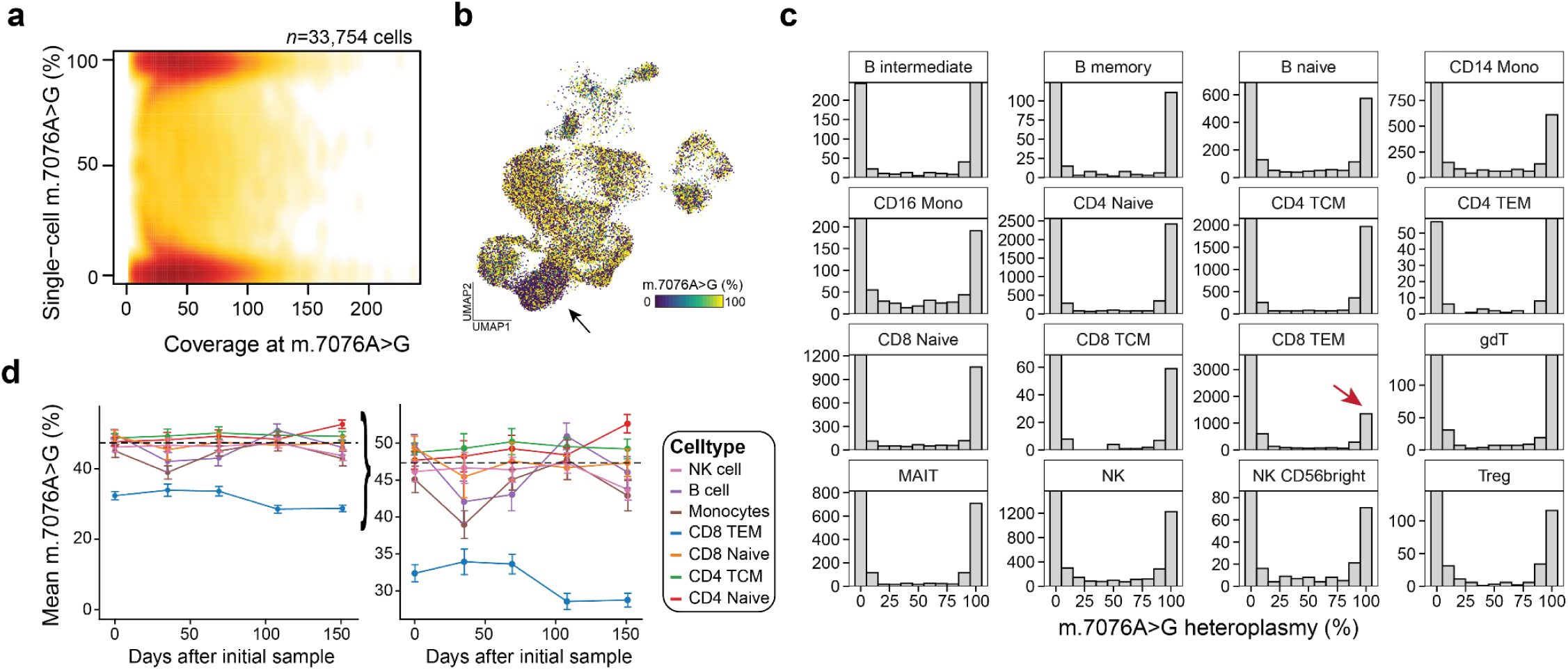
Supporting information for selection against mutant m.7076G in CD8^+^ TEM. **(a)** Heatmap of single-cells comparing the coverage of both m.7076 alleles against heteroplasmy. **(b)** Compare to Fig. 1e with the smoothed representation. **(c)** Histograms comparing the 16 most common cell types from the Azimuth/Bridge Integration annotation. The red arrow highlights the significant reduction of cells with the m.7076A>G variant specifically in CD8^+^ TEM cells, but not other cell types. **(d)** Longitudinal heteroplasmy of different cell populations over >150 days of sampling. The right panel is a zoom of the region on the left panel (note axis). Pseudobulk heteroplasmy estimates including the standard error of the mean are shown.

**Extended Data Fig. 2.**
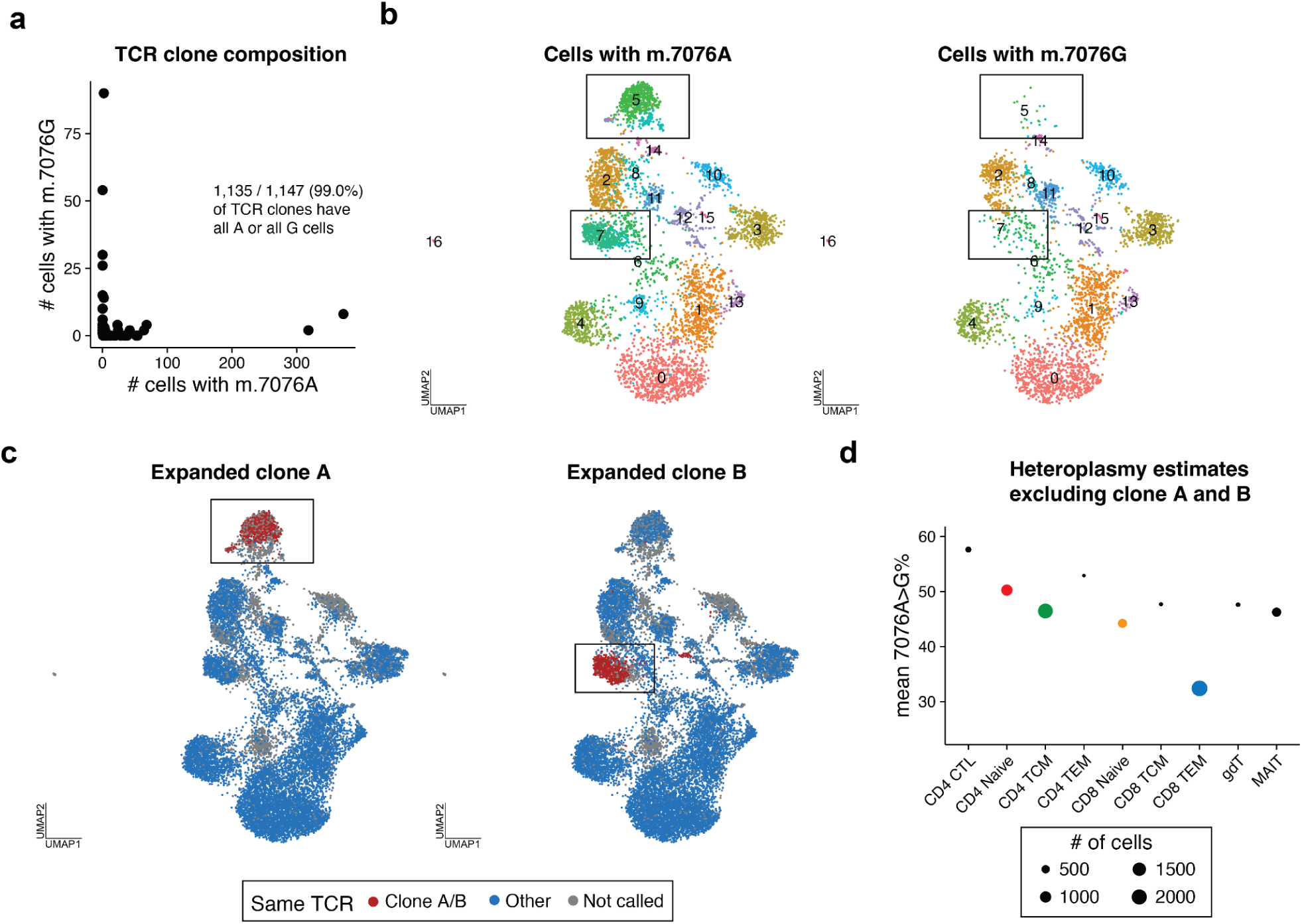
Supporting information for selection against mutant m.7076A>G using scRNA-seq. **(a)** Overlap of TCR clones with respective m.7076 alleles. Each dot is a TCR clone (min. 2 cells) summarizing the number of cells with either wildtype m.7076A or mutant m.7076G homoplasmy. **(b** Unsupervised clustering and dimensionality reduction of T cells stratified by m.7076A or m.7076G homoplasmy. Black boxes around clusters 5 (98.1%) and 7 (95.9%) represent specific cell states that are primarily restricted to cells harboring the m.7076A allele. **(c)** Annotation of two highly expanded TCR clones restricted to cells homoplasmic for wildtype m.7076A. Blue represents TCR clones (n≥2 cells) that were not highly expanded. **(d)** Heteroplasmy of the m.7076A>G allele in indicated T cell subpopulations based on scRNA-seq after excluding clones A and B from panel (c). The size of each dot is scaled by the abundance of cells in each cell state.

**Extended Data Figure 3.**
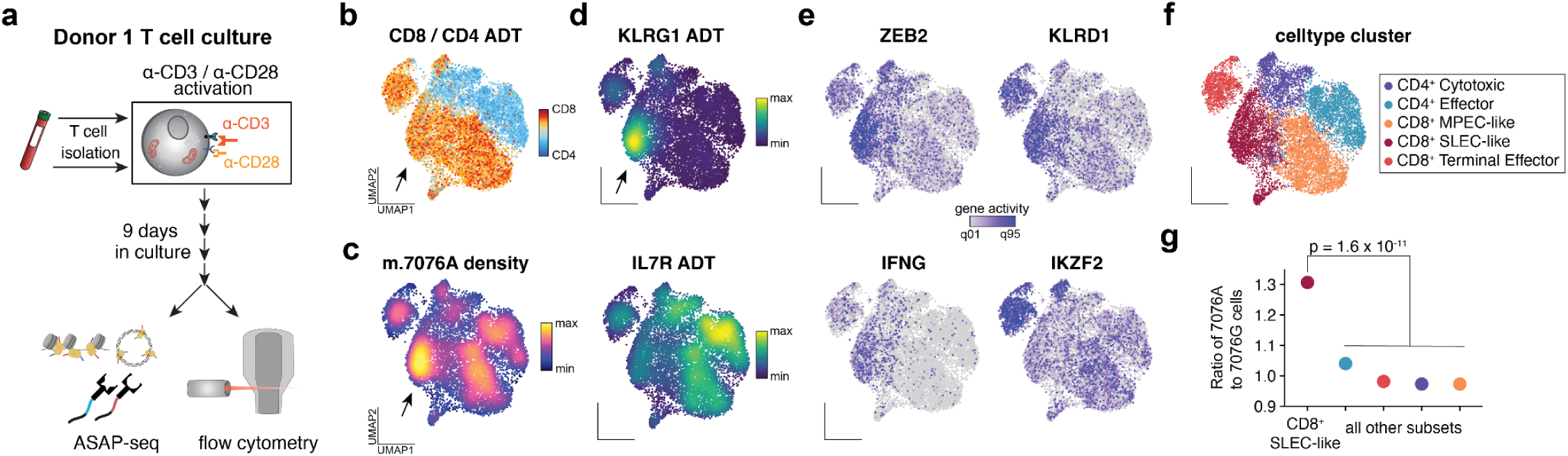
*In vitro* activation of T cells refines cell states depleted of the mutant 7076G allele. **(a)** Schematic of experimental design. T cells were isolated from Donor 1, *in vitro* activated, and cultured for 9 days before profiling via flow cytometry and ASAP-seq. **(b)** UMAP of accessible chromatin profiles and projected ratio of CD8 over CD4 antibody-derived tags from day 9 cells profiled via ASAP-seq. Arrow indicates a population highly enriched for the m.7076A (wildtype) allele. **(c)** Same as (b) but colored by the density of the m.7076A (wildtype) allele. **(d)** UMAP embedding colored by KLRG1 (top) and IL7R (bottom) antibody tag density. **(e)** UMAP colored by selected gene activity scores for four indicated gene loci. **(f)** UMAP colored by indicated cell state cluster. **(g)** Ratio of wildtype m.7076A to mutant m.7076G cells within indicated cell states. P-value represents the statistical significance of a two-sided binomial test statistic.

**Extended Data Fig. 4.**
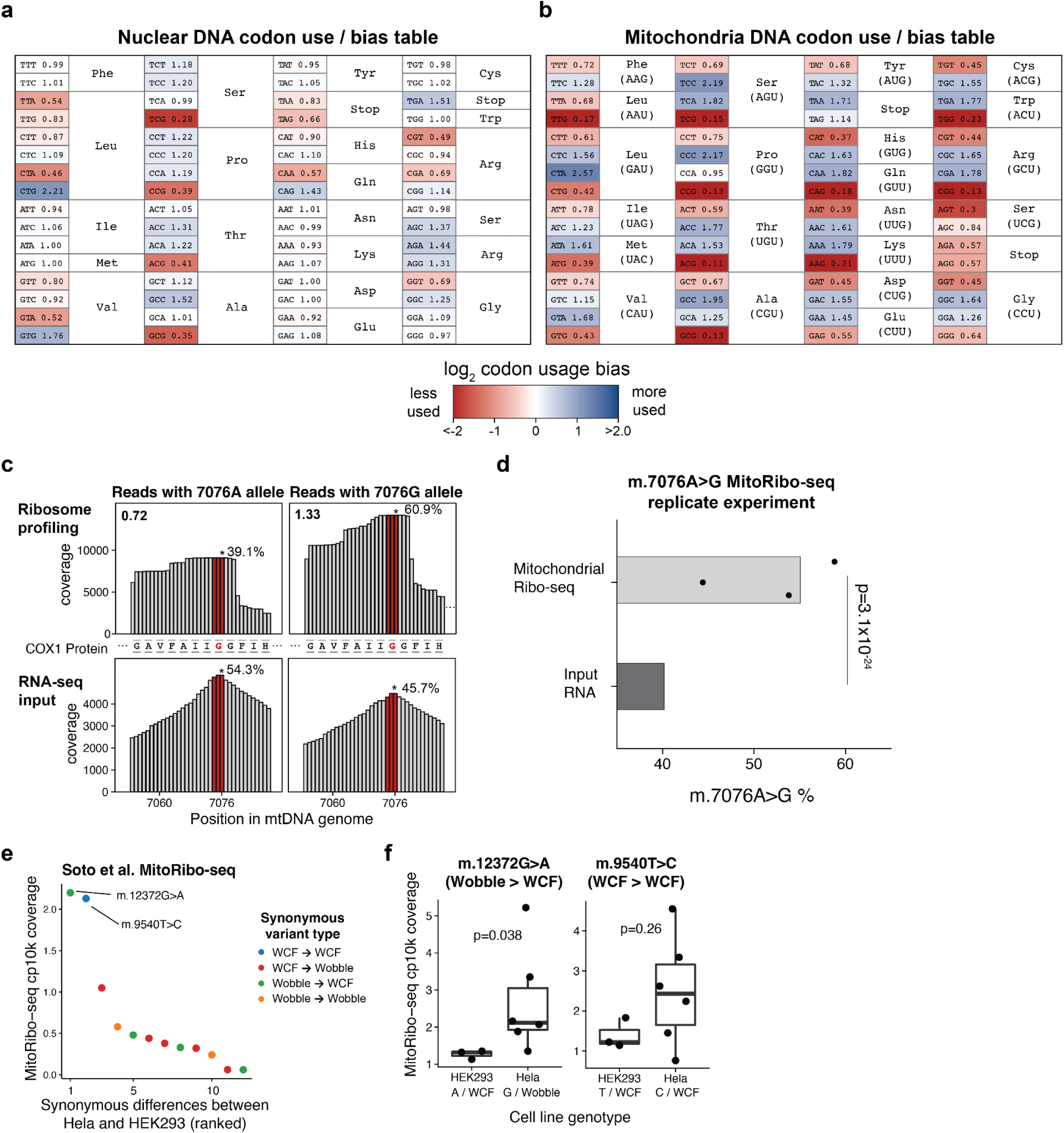
Supporting information for analysis of the mitochondrial tRNA pool and impact on translational efficiency via the wobble effect. **(a)** Codon bias table for the nuclear genome. Within each amino acid, the ratio of observed codon usage over a null model of equal codon use per amino acid and colored by the log_2_ of this measure is shown. **(b)** Same as (a) but for the 13 polypeptides encoded in the mitochondrial genome. **(c)** Coverage near the m.7076A>G variant. Red bars indicate the mutated codon with the m.7076 allele (noted with an asterisk). The relative proportion of reads phased to either allele per library is indicated. The translation pause ratio, defined as the fraction of reads from ribosome profiling over the RNA-seq libraries, is noted in the top left corner. **(d)** Three additional replicate fractions of MitoRibo-seq from an independent experiment of Donor 1 cells. Statistical significance was determined using a Fisher’s exact test of 7076A and G alleles summed between replicates. **(e)** Synonymous alleles with distinct homoplasmy between HEK293 and Hela cell lines from a previous study^23^. Two variants called out had sufficient coverage (>2 counts per 10k; cp10k) for further analysis. **(f)** Comparison of MitoRibo-seq read abundances between cell lines for two variants highlighted in (e). Shown are two variants that differ between cell lines, including one that impacted wobble-dependent translation (m.12372G>A; Wobble→WCF; increased stalling at wobble allele) or not (m.9540T>C; WCF→WCF). Statistical test: two-sided Wilcoxon test.

**Extended Data Fig. 5.**
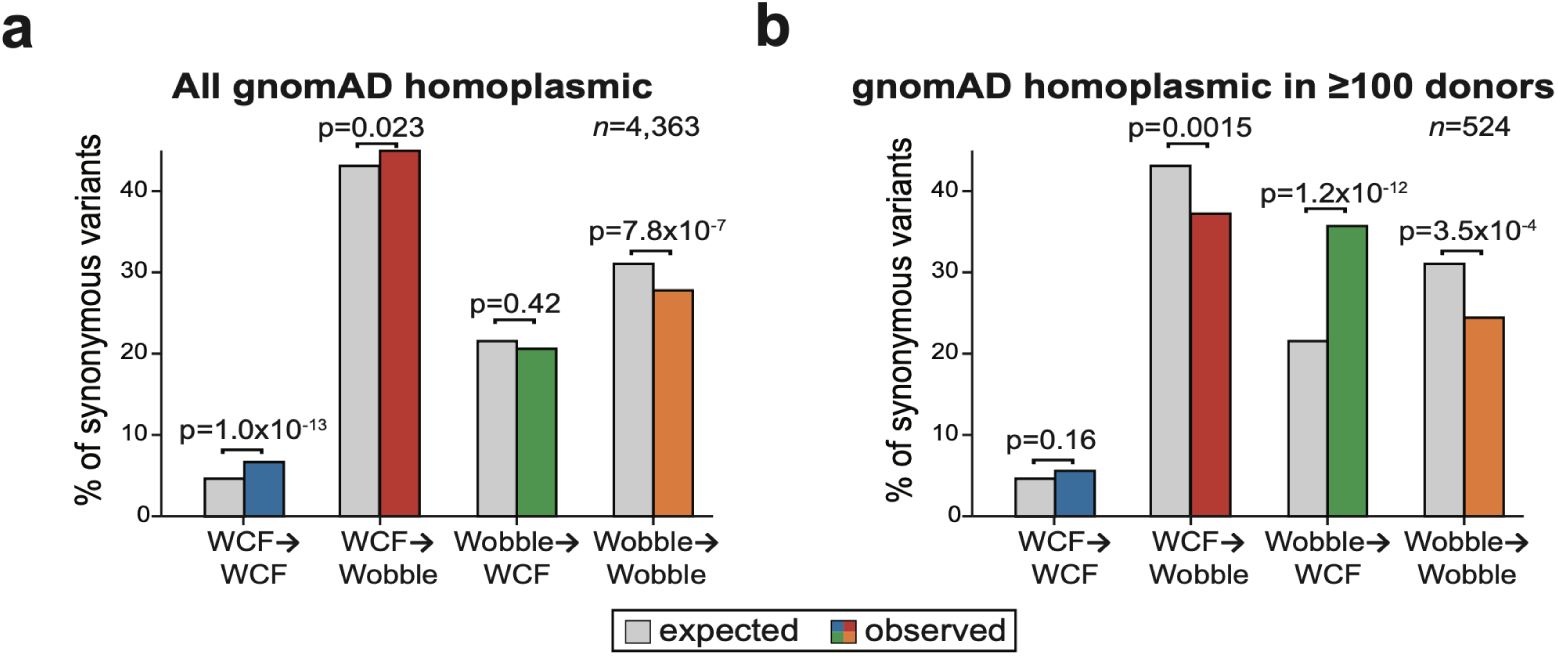
Supporting information comparing mtDNA synonymous mutation patterns in gnomAD sequencing data. **(a)** Comparison of % observed (color) versus expected (grey) synonymous mtDNA mutations from gnomAD for homoplasmic variants (all observed homoplasmic variants from the population). **(b)** Same as in (a) but for variants present in ≥100 healthy individuals based on gnomAD analysis, showing enrichment of more optimal Wobble → WCF synonymous codons. P-values represent the statistical significance of a two-sided binomial test statistic.

**Extended Data Fig. 6.**
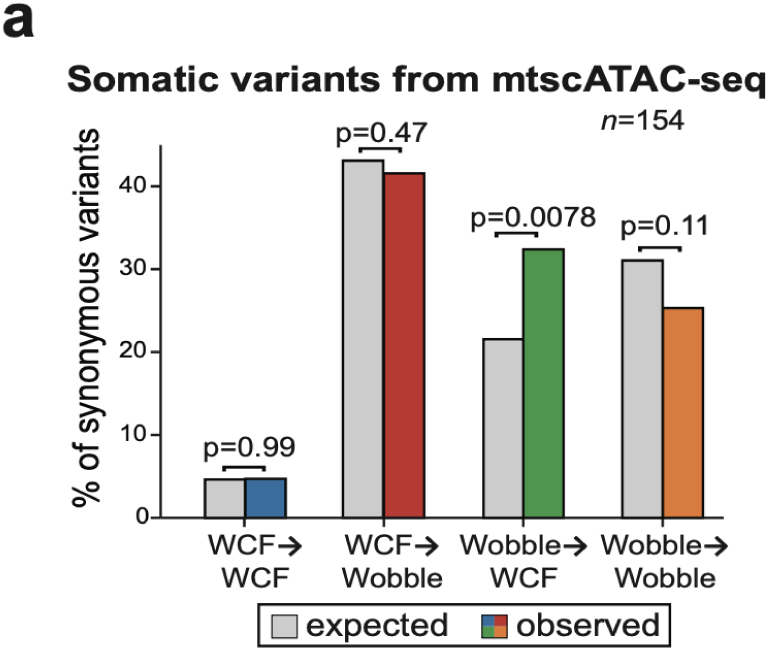
Supporting information comparing mtDNA synonymous mutation patterns in healthy human tissues. **(a)** Comparison of % observed (color) versus expected (grey) synonymous mtDNA mutations from a healthy 47-year-old individual profiled with mtscATAC-seq for somatic heteroplasmic variants in PBMCs.

**Extended Data Figure 7.**
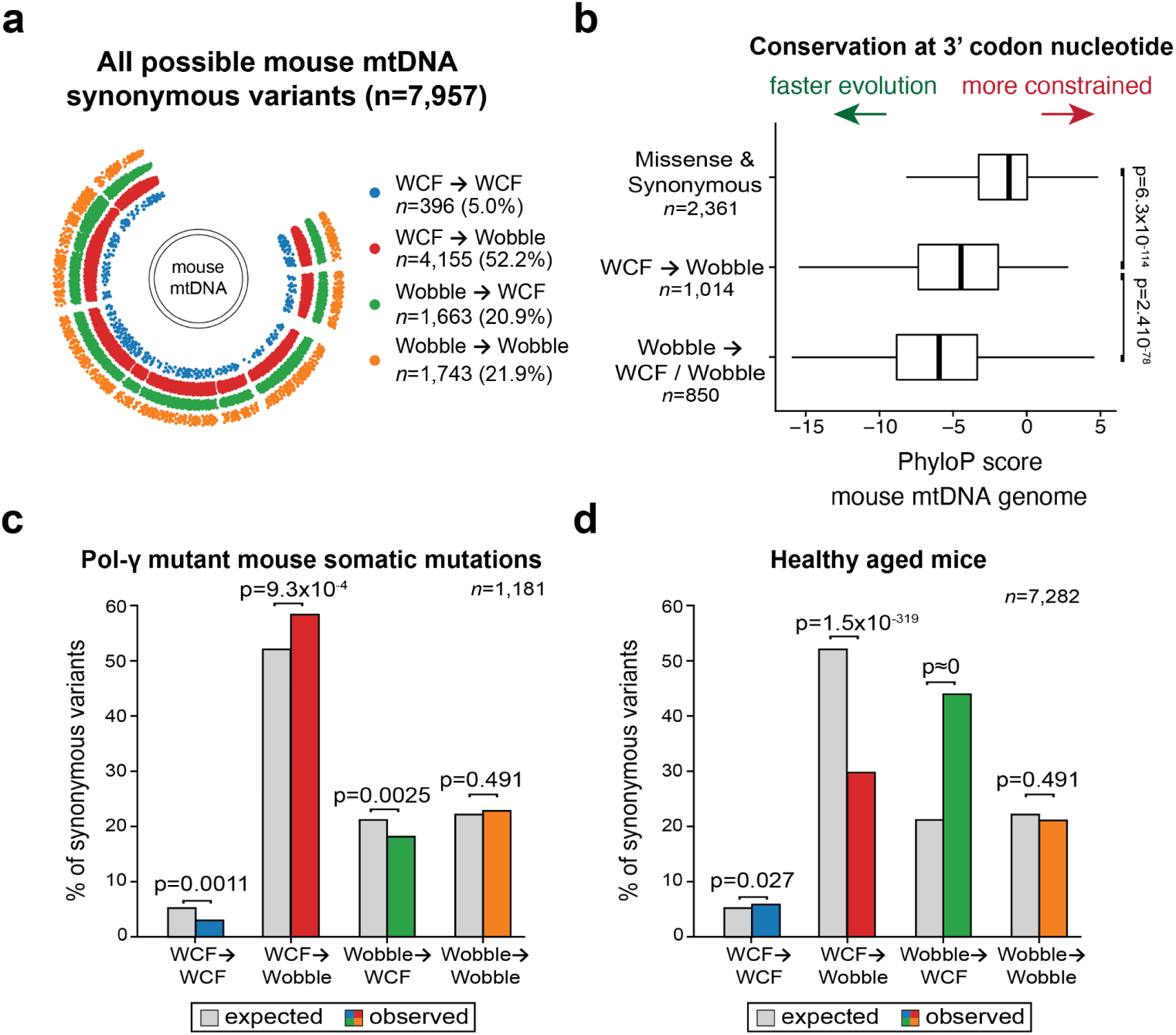
A new ontogeny of synonymous mtDNA mutations in the murine genome via wobble-dependent translation. **(a)** Classification of all 7,957 possible synonymous mtDNA variants based on Watson-Crick-Franklin (WCF) or wobble-dependent base-pairing at either the reference or alternative allele. **(b)** Inter-species conservation at wobble-position nucleotides in mitochondrial codons for the murine genome. Reference alleles that can be mutated to different outcomes are specified and grouped with the number of wobble-position codons in each class noted. P-values represent a Wilcoxon test. **(c)** Comparison of observed (color) versus expected (grey) heteroplasmic synonymous mtDNA mutations occurring in the pol-γ mutant mice^36^ and **(d)** in healthy aging mice^37^. P-values represent the statistical significance of a two-sided binomial test statistic.

## Methods

### Mitochondrial single-cell ATAC-seq (mtscATAC-seq)

MtscATAC-seq libraries were generated using the 10x Chromium Controller and the Chromium Single Cell ATAC Library & Gel Bead Kit (#1000175) according to the manufacturer’s instructions (CG000209-Rev G; CG000168-Rev B) as outlined below and previously described to increase mtDNA yield and genome coverage^9^. Briefly, 1.5 ml or 2 ml DNA LoBind tubes (Eppendorf) were used to wash cells in PBS and downstream processing steps. After washing cells were fixed in 1% formaldehyde (FA; ThermoFisher #28906) in PBS for 10 min at RT, quenched with glycine solution to a final concentration of 0.125 M before washing cells twice in PBS via centrifugation at 400 g, 5 min, 4°C. Cells were subsequently treated with lysis buffer (10mM Tris-HCL pH 7.4, 10mM NaCl, 3mM MgCl_2_, 0.1% NP40, 1% BSA) for 3 min for primary cells on ice, followed by adding 1 ml of chilled wash buffer and inversion (10mM Tris-HCL pH 7.4, 10mM NaCl, 3mM MgCl_2_, 1% BSA) before centrifugation at 500 g, 5 min, 4°C. The supernatant was discarded and cells were diluted in 1x Diluted Nuclei buffer (10x Genomics) before counting using Trypan Blue and a Countess II FL Automated Cell Counter. If large cell clumps were observed a 40 µm Flowmi cell strainer was used before processing cells according to the Chromium Single Cell ATAC Solution user guide with no additional modifications. Briefly, after tagmentation, the cells were loaded on a Chromium controller Single-Cell Instrument to generate single-cell Gel Bead-In-Emulsions (GEMs) followed by linear PCR as described in the protocol using a C1000 Touch Thermal cycler with 96-Deep Well Reaction Module (BioRad). After breaking the GEMs, the barcoded tagmented DNA was purified and further amplified to enable sample indexing and enrichment of scATAC-seq libraries. The final libraries were quantified using a Qubit dsDNA HS Assay kit (Invitrogen) and a High Sensitivity DNA chip run on a Bioanalyzer 2100 system (Agilent).

### Single-cell RNA-seq and TCR profiling

Libraries for scRNA-seq were generated using the 10x Chromium Controller and the Chromium Single Cell 5′ Library Construction Kit and human B cell and T cell V(D)J enrichment kit according to the manufacturer’s instructions. Briefly, the suspended cells were loaded on a Chromium controller Single-Cell Instrument to generate single-cell Gel Bead-In-Emulsions (GEMs) followed by reverse transcription and sample indexing using a C1000 Touch Thermal cycler with 96-Deep Well Reaction Module (BioRad). After breaking the GEMs, the barcoded cDNA was purified and amplified, followed by fragmenting, A-tailing, and ligation with adaptors. Finally, PCR amplification was performed to enable sample indexing and enrichment of scRNA-seq libraries. For T-cell receptor sequencing, target enrichment from cDNA was conducted according to the manufacturer’s instructions. The final libraries were quantified using a Qubit dsDNA HS Assay kit (Invitrogen) and a High Sensitivity DNA chip run on a Bioanalyzer 2100 system (Agilent). 10x scRNA-seq libraries were sequenced as recommended by the manufacturer (∼20,000 reads per cell) via a NovaSeq 6000 using an S4 flow cell.

### ATAC with Selected Antigen profiling by sequencing (ASAP-seq)

Cultured primary T cells were stained with a TSA-conjugated antibody panel (BioLegend ‘Universal’ Totalseq-A panel) that targets 154 distinct epitopes as previously described^11^. Briefly, following sorting, cells were fixed in 1% formaldehyde and processed as described for the mtscATAC-seq workflow described above, with the modification that during the barcoding reaction, 0.5µl of 1µM bridge oligo A (BOA for TSA) was added to the barcoding mix. For GEM incubation the standard thermocycler conditions were used as described by 10x Genomics for scATAC-seq. Silane bead elution and SPRI cleanup steps were modified as described to generate the indexed protein tag library^11^. The final libraries were quantified using a Qubit dsDNA HS Assay kit (Invitrogen) and a High Sensitivity DNA chip run on a Bioanalyzer 2100 system (Agilent). Libraries were amplified, sequenced, and preprocessed as previously described^11^.

### Single-cell ATAC-seq analyses

Raw sequencing data were demultiplexed using CellRanger-ATAC mkfastq. Demultiplexed sequencing reads for all libraries were aligned to the mtDNA blacklist modified^9^ hg38 reference genome using CellRanger-ATAC count v2.0. Mitochondrial DNA genotypes were determined using the mgatk workflow with default parameters^9^. Cell state analyses, including gene activity scores and surface protein visualization, were performed using the Seurat/Signac framework^49, 50^. For PBMC cell type annotations, granular cell type labels and UMAP coordinates were established by using the Seurat Dictionary Learning^51^ for cross-modality integration. We used Azimuth CITE-seq reference dataset labels^52^ with public 10x Genomics Multiome RNA- and ATAC-seq PBMC data as a cross-modality bridge. To assess whether additional genetic chimerism was present in cells marked with m.7076A>G, we split the full single-cell .bam file based on high-confidence cells with either the m.7076A or m.7076G allele and used FreeBayes^53^ to genotype variants in the nuclear chromosomes using pseudobulk .bam files of the scATAC-seq profiles. After intersecting the two .vcf files from either the m.7076A or m.7076G variant calls, we did not identify evidence of high-quality variants (QUAL>100) specific to either allele, noting that our variant calling was restricted to accessible chromatin regions.

### Single-cell RNA-seq analyses

Raw sequencing data were demultiplexed using CellRanger mkfastq and aligned to the host reference genome using CellRanger v6.0 and TCR sequences were processed using the CellRanger vdj pipeline with default settings. Mitochondrial DNA genotypes were determined using the mgatk workflow with default parameters^9^ while accounting for UMIs using the -ub tag. T cell subset annotations were derived using the Azimuth CITE-seq reference dataset labels^52^ from healthy 10x PBMC data. Cells were assigned as m.7076A or m.7076G using a minimum coverage of 2 UMIs and homoplasmy for either allele from the transcriptomic data (mean coverage = 3.2x/cell from 5’ scRNA-seq for m.7076). Expression levels of *MT-CO1* were determined from the default normalization in Seurat and assessed for differences using a Wilcoxon test with the Benjamini-Hochberg adjustment for multiple testing.

### Mitochondrial ribosome profiling

Peripheral blood mononuclear cells were cultured in RPMI-1640 medium supplemented with 10% Fetal Bovine Serum (FBS), penicillin and streptomycin, and 10 ng/ml IL-2 (PeproTech) at 37°C, 5% CO_2_. For *in vitro* expansion, cells were stimulated with Dynabeads Human T-Activator αCD3/αCD28 at a bead-to-cell ratio of 1:2 (11131D, Thermo Fisher Scientific). Cell counts were determined every 2-3 days, and maintained at a density of 1-2×10^6^ cells/ml. On day 7 after stimulation, cells were collected for mitochondrial ribosome profiling (MitoRiboSeq). Briefly, 0.1 mg/ml cycloheximide (CHX) and 0.1 mg/ml chloramphenicol (CAP) were directly added into the cell culture media followed by 5 min incubation at 37°C, 5% CO_2_. 20×10^6^ cells were then centrifuged and washed with ice-cold PBS supplied with 0.1 mg/ml CHX and 0.1 mg/ml CAP. Dynabeads were magnetically removed, cells were pelleted, snap-frozen in liquid nitrogen, and stored at -80°C until further processing.

The protocol for Mitochondrial ribosome enrichment was adapted from previous work^54^. The flash-frozen cell pellet was resuspended in 1 mL of lysis buffer (20 mM Tris-HCl pH 8.0, 100 mM KCl, 10 mM MgCl2, 0.1 mg/mL chloramphenicol, 0.1 mg/mL cycloheximide, 1 mM DTT, 1% (vol/vol) Triton X-100, 0.1% (vol/vol) NP-40, 1× complete protease inhibitor cocktail, 20 U/mL RNasin, 20 U/mL DNase I) and incubated for 10 min on ice. The lysate was homogenized by passing it 3 times through a 26 G needle. Cell debris was removed by centrifugation at 16,000g for 10 min (4°C) and 100 µL of clarified lysate was kept aside for preparation of RNA-seq libraries. Ribosome-protected fragments (RPFs) were generated by nuclease digestion. To this end, 3.75 U/µL of micrococcal nuclease (NEB) and 5 mM CaCl2 were added to the remaining cell lysate; the incubation was performed at room temperature with gentle shaking for 1h. The reaction was quenched by adding 6 mM of EGTA. The digested lysate was loaded on top of a linear 5-45% sucrose gradient containing 20 mM Tris-HCl pH 8.0, 100 mM KCl, 10 mM MgCl2, 0.1 mg/mL chloramphenicol, 0.1 mg/mL cycloheximide and 1 mM DTT and centrifuged at 35,000 rpm, 4°C for 2 hr using a Beckmann SW40 rotor. The gradient was then fractionated into 20 fractions of 0.57 mL using a Biocomp gradient station fractionator, which allowed the recording of a UV absorbance profile at 254 nm.

### Western blot of gradient fractions

Western blot analysis was used to monitor the successful isolation of the 55S ribosome. Proteins were precipitated from 230 µL of each gradient fraction and 25 µL of input lysate by adding Trichloroacetic acid (TCA) to a final concentration of 20 %. Samples were incubated on ice for 1-2 h and proteins were precipitated by centrifugation at 15,000g at 4°C. The supernatant was removed and protein pellets were washed twice with 300 µL ice-cold acetone, followed by a 15 min centrifuging at 15,000g and 4°C. Pellets were air-dried and resuspended in 30 fL 1x Lämmli-Buffer. Samples were separated on an 8 12% Bis-Tris SDS-PAGE gel and transferred to a nitrocellulose membrane using the iBlot/iBind system (Thermo Fisher Scientific). The presence of mitochondrial and cytoplasmic ribosomes in gradient fractions was estimated by incubating resulting membranes with anti-MRPL11 monoclonal antibody (D68F2, Cell Signaling) and anti-RPS6 rabbit monoclonal antibody (Cell Signaling).

### RPF isolation and library generation

Ribosome-protected fragments were isolated using Phenol/Chloroform extraction. To that end, 300 µl of the corresponding gradient fractions were mixed with equal amounts of Phenol:Chloroform:IAA (25:24:1, pH 6.6), transferred to a PhaseLock tube (VWR International), and centrifuged at 15,000g for 5 min. A second clean-up step was performed using the isolated aqueous phase and adding equal amounts of Chloroform. After centrifugation, the aqueous phase was isolated and purified using the RNA Clean & Concentrator kit (Zymo Research) by following the manufacturer’s instructions for small RNAs. Subsequent library preparation was performed as previously described^55^ with the following modification: ribosomal RNA depletion was skipped for RPF libraries due to low starting material. For the preparation of matched RNA-seq libraries, total RNA was extracted from 40 µL of clarified input lysate using the Direct-zol RNA MiniPrep Kit (Zymo Research). Subsequently, rRNA was depleted using riboPOOLs (siTOOLs Biotech) and following the manufacturer’s instructions. Subsequently, RNA-seq libraries were generated as described previously^55^.

### MitoRibo-seq Analyses

Raw .fastq files were trimmed for adapter sequences using cutadapt and subsequently aligned with bwa mem using default parameters as previously described^55^. Summary statistics of coverage and m.7076A>G heteroplasmy were determined using the mgatk workflow with default parameters as well as the -kd flag to retain duplicate fragments with the same start and end coordinates. Pileup coverages of the locus split by the m.7076 allele were determined using a custom python script using pysam library.

To examine other variants aside from m.7076A>G that may participate in wobble-mediated translational stalling, we utilized high-quality HEK293 (n=3) and Hela (n=6) MitoRibo-seq profiles from a recent manuscript^23^ at GEO accession GSE173283. Raw RNA-seq reads were processed and variants distinct between the two cell lines were inferred using mgatk^9^. We identified two synonymous variants that were homoplasmic but with distinct alleles in the two cell lines that had mean counts per 10,000 reads (cp10k) greater than 2 in the MitoRibo-seq data.

### Annotation of synonymous variation in the mitochondrial genome

The revised Cambridge reference sequence (used in GRCh37, GRCh38, and hg38) was used as the basis for the ontogeny of somatic variation. We utilized all possible mtDNA variations and protein-coding annotations as previously described^10^. Using this landscape of 8,284 synonymous mutations, we annotated whether the codon in the reference or the alternate allele would create a canonical (i.e., Watson-Crick-Franklin, WCF) base-pairing between the codon and anticodon or would require wobble-dependent base-pairing for translation. Once these annotations were established, the null model of the abundance of each variant class (**Fig. 4**) was derived from the empirical distribution of these variant classes in the full mitochondrial genome. To compare population-level enrichment, we used the gnomAD v3.1 variant call set via the MT .vcf file available on the download page^24^ and parsed the HelixMTdb database available online. For the analysis of somatic variation in cancer samples (**Fig. 6**), we utilized three pan-cancer datasets of somatic mtDNA variation derived from tumors that were previously reported^34^. For the overall analysis of somatic cancer mutations (colored bars), we concatenated the variant lists, which effectively provided a weighted average proportional to the number of mutations called per study. GTEx analyses of somatic variants were determined using a previous catalog that filtered somatic variants occurring in more than one BioSample to reduce potential biases from variant calling in transcriptomics data^13^.

To further separate synonymous mtDNA variants based on tRNA class (**Fig. 6b**), we categorized each of the 22 human mt-tRNAs based on one of five categories using the chemical RNA structures previously annotated^33^. These classes were determined based on the chemical structure of the base in the wobble position anticodon (position 34 in the tRNA structure). Four of the five classes and chemical structures are shown in **Fig. 6b** with the fifth being the f_5_C34 which is restricted to the methionine codon, which we excluded from our analysis as it was only one tRNA and two total codons. Otherwise, each variant was annotated with the tRNA responsible for translation based on the canonical mtRNA translation table^33^.

### Evolutionary and cross-species analyses

DNA variants from the mitochondrial genome used to define haplogroups were determined from the PhyloTree Build 17 annotation as distributed by HaploGrep2^26, 27^. Filtered variants required a phylogenetic recurrence of 10, leading to 289 variants, including 48 missense and 87 synonymous variants. To calculate the inter-species evolution of the mitochondrial genome, we utilized the phyloP annotation per nucleotide (rather than per variant and this metric is not available from phyloP), requiring our annotation of wobble-position nucleotides in the mitochondrial genome to be assigned to one of three categories (**Fig. 4b**). The associations shown in **Fig. 4b** are for a 20-species phyloP calculation but were consistent using a 100-species estimation as well. Both phyloP annotations were downloaded from UCSC Genome Browser for the GRCh38 genome annotation. Similar analyses for the murine mm10 genome, including a 60-species phyloP, are shown in **Extended Data Fig. 7**. Somatic variants from the pol-γ mutant mice^36^ and in healthy aging mice^37^ were downloaded from the supplemental tables from prior publications.

### Complex human trait analysis

#### Cohort and Phenotypes

The UK Biobank is a prospective study of approximately 500,000 participants 40–69 years of age at recruitment. Participants were recruited in the UK between 2006 and 2010 and are continuously followed. The average age at recruitment for sequenced individuals was 56.5 years. Participant data include health records that are periodically updated by the UKB, self-reported survey information, linkage to death and cancer registries, collection of urine and blood biomarkers, imaging data, accelerometer data, genetic data, and various other phenotypic endpoints. All study participants provided informed consent.

We harmonized the UKB phenotype data as previously described. Briefly, we studied two main phenotypic categories: binary and quantitative traits taken from the December 2021 data release (UKB application 26041). We parsed phenotypic data using our previously described R package, PEACOK (https://github.com/astrazeneca-cgr-publications/PEACOCK). In addition, as previously described, we grouped relevant ICD-10 and ICD-9 codes into clinically meaningful “Union” phenotypes as previously described^56^. For all binary phenotypes, we matched controls by sex when the percentage of female cases was significantly different (Fisher’s exact two-sided P < 0.05) from the percentage of available female controls. In total, we tested for associations with 18,688 binary phenotypes and 1,656 quantitative phenotypes.

#### Whole-genome sequencing

Whole-genome sequencing (WGS) data of the UKB participants were generated by deCODE Genetics and the Wellcome Trust Sanger Institute as part of a public-private partnership involving AstraZeneca, Amgen, GlaxoSmithKline, Johnson & Johnson, Wellcome Trust Sanger, UK Research and Innovation, and the UKB. These individuals were pseudorandomly selected from the set of UKB participants. The WGS sequencing methods have been previously described. Briefly, genomic DNA underwent paired-end sequencing on Illumina NovaSeq6000 instruments with a read length of 2×151 and an average coverage of 32.5x. Conversion of sequencing data in BCL format to FASTQ format and the assignments of paired-end sequence reads to samples were based on 10-base barcodes, using bcl2fastq v2.19.0. Initial quality control was performed by deCODE and Wellcome Sanger, which included sex discordance, contamination, unresolved duplicate sequences, and discordance with microarray genotyping data checks. A total of 199,949 genomes passed these quality control measures.

UK Biobank genomes were processed at AstraZeneca using the provided CRAM format files. A custom-built Amazon Web Services (AWS) cloud compute platform running Illumina DRAGEN Bio-IT Platform Germline Pipeline v3.7.8 was used to align the reads to the GRCh38 genome reference and to call small variants, including on the mitochondrial genome where a continuous allele frequency model is used; a single alternate allele is considered as a candidate variant and an allele fraction is estimated for emitted variants. All PASS variants emitted had a confidence score (LOD) above the default of 6.3. Mitochondrial SNVs and indels were annotated using SnpEff v4.3^57^ against Ensembl Build 38.92^58^ with the use of the vertebrate mitochondrial amino acid code configured using ‘hg38.M.codonTable: Vertebrate_Mitochondrial’ and ‘hg38.MT.codonTable: Vertebrate_Mitochondrial’.

From an initial cohort of 199,949 genomes, we removed 120 where the genome showed kinship<0.49 (<98% identity) to the exome from the same participant, where available; 8 that could not be linked to a participant; 0 with ≥4% contamination estimated by VerifyBamID^59^; 5 where self-reported gender did not match karyotypic sex inferred by X:Y coverage ratio across CCDS release 22 bases; 0 where <94.5% of CCDS r22 bases had ≥10-fold coverage; 100 in the top 0.05% of CCDS r22 coverage (>67.2x); and 4,915 to obtain a set with all pairwise kinship estimates ≤0.1769 based on KING.

Europeans are the most well-represented genetic ancestry in the UKB. We identified the participants with European genetic ancestry based on Peddy^60^ v0.4.2 Pr(EUR)>0.95. We then performed finer-scale ancestry pruning of these individuals, retaining those within four standard deviations from the mean across the first four principal components, resulting in a final set of 183,116 not closely related genomes of European ancestry for analysis.

#### Variant-level association tests

Considering all 8,284 possible synonymous variants, we performed variant-level association tests for the 2,786 synonymous variants by requiring the alternate allele to be observed in at least six individuals of European ancestry. We used a two-sided Fisher’s exact test for binary traits and linear regression for quantitative traits (correcting for age and sex). For binary traits, we only included phenotypes with at least 30 cases. We dichotomized the alternate allele read fraction into a binary genotype indicator variable, with fractions ≥0.90 coded as 1 and fractions ≤0.10 (including the absence of a variant call) coded as zero. Intermediate fractions were set to missing genotypes. Variants were required to have a mapping quality score (MQ) ≥ 40 and DRAGEN variant status PASS, i.e. not filtered with lod_fstar or base_quality.

We performed an n-of-1 permutation test, as previously described in our exome-based phenome-wide association study, to determine an appropriate p-value threshold. Briefly, we shuffled the case-control (or quantitative measurement) labels once for every phenotype while maintaining the participant-genotype structure. At a p-value threshold of 1×10^-6^, a total of 8/52,064,768 binary and 1/4,613,616 associations in the permuted analysis were significant, suggesting a negligible false positive rate for the thresholds reported in the main text, and thus used as a conservative threshold for analysis and interpretation.

## Acknowledgments

We thank members of the Satpathy lab for helpful discussions. CAL, LSL, and ATS are supported by the National Institutes of Health grant UM1HG012076. C.A.L. is supported by a Stanford Science Fellowship, a Parker Institute for Cancer Immunotherapy Scholarship, and NHGRI K99HG012579. A.T.S. is supported by the Burroughs Wellcome Fund Career Award for Medical Scientists, the Parker Institute for Cancer Immunotherapy, a Pew-Stewart Scholars for Cancer Research Award, a Cancer Research Institute Lloyd J. Old STAR Award, a Scholar Award from the American Society of Hematology, and a Baxter Foundation Faculty Scholar Award. YHH is supported by a PhD fellowship from the Hector Fellow Academy. PK and LN receive support as associate members of the Hector Fellow Academy. LSL is supported by an Emmy Noether fellowship by the German Research Foundation (DFG, LU 2336/2-1), a Longevity Impetus grant, and a Hector Research Career Development Award by the Hector Fellow Academy.

## Author contributions

C.A.L., L.S.L., and A.T.S. conceived and designed this work. C.A.L. lead all analyses and is responsible for the accuracy and reproducibility of the manuscript. Y.Y., A.S.G., T.A., S.Z., R.R.S., K.S., and B.D. performed genomics experiments. J.C.G., V.L., J.C.U., R.S., W.J.G., and A.K. contributed to data analysis and interpretation. R.S.D., Q.Q., F.H., K.R.S., S.V.V.D., and S.P. performed the human genetics analyses. Y.H.H., L.N., F.A.B., F.W., P.Y., Z.M. performed or advised on T cell experiments. C.A.L., M.M., L.S.L., and A.T.S. supervised the work. C.A.L., L.S.L., and A.T.S. wrote the manuscript with input from all authors.

## Code and Data Availability

Sequencing data associated with this work is available at GEO accession **GSE216915** with reviewer access **ijmrugiwflqzdif**. All custom code to reproduce all analyses supporting this manuscript is available at https://github.com/caleblareau/7076.

## References

1. Walker, M. A. et al. Purifying Selection against Pathogenic Mitochondrial DNA in Human T Cells. N. Engl. J. Med. 383, 1556–1563 (2020).

2. Lareau, C. A. et al. Single-cell multi-omics reveals dynamics of purifying selection of pathogenic mitochondrial DNA across human immune cells. Preprint at https://doi.org/10.1101/2022.11.20.517242.

3. Khajuria, R. K. et al. Ribosome Levels Selectively Regulate Translation and Lineage Commitment in Human Hematopoiesis. Cell 173, 90–103.e19 (2018).

4. Stewart, J. B. & Chinnery, P. F. The dynamics of mitochondrial DNA heteroplasmy: implications for human health and disease. Nat. Rev. Genet. 16, 530–542 (2015).

5. Yazar, S. et al. Single-cell eQTL mapping identifies cell type–specific genetic control of autoimmune disease. Science 376, eabf3041 (2022).

6. Nathan, A. et al. Single-cell eQTL models reveal dynamic T cell state dependence of disease loci. Nature 606, 120–128 (2022).

7. Nam, A. S. et al. Single-cell multi-omics of human clonal hematopoiesis reveals that DNMT3A R882 mutations perturb early progenitor states through selective hypomethylation. Nat. Genet. 54, 1514–1526 (2022).

8. Nam, A. S. et al. Somatic mutations and cell identity linked by Genotyping of Transcriptomes. Nature 571, 355–360 (2019).

9. Lareau, C. A. et al. Massively parallel single-cell mitochondrial DNA genotyping and chromatin profiling. Nat. Biotechnol. (2020) doi:10.1038/s41587-020-0645-6.

10. Miller, T. E. et al. Mitochondrial variant enrichment from high-throughput single-cell RNA sequencing resolves clonal populations. Nat. Biotechnol. 40, 1030–1034 (2022).

11. Mimitou, E. P. et al. Scalable, multimodal profiling of chromatin accessibility, gene expression and protein levels in single cells. Nat. Biotechnol. (2021) doi:10.1038/s41587-021-00927-2.

12. Fiskin, E. et al. Single-cell profiling of proteins and chromatin accessibility using PHAGE-ATAC. Nat. Biotechnol. (2021) doi:10.1038/s41587-021-01065-5.

13. Ludwig, L. S. et al. Lineage Tracing in Humans Enabled by Mitochondrial Mutations and Single-Cell Genomics. Cell 176, 1325–1339.e22 (2019).

14. Bolze, A. et al. A catalog of homoplasmic and heteroplasmic mitochondrial DNA variants in humans. Preprint at https://doi.org/10.1101/798264.

15. Karczewski, K. J., Francioli, L. C. & MacArthur, D. G. The mutational constraint spectrum quantified from variation in 141,456 humans. Yearbook of Paediatric Endocrinology Preprint at https://doi.org/10.1530/ey.17.14.3 (2020).

16. Jones, N. et al. Metabolic Adaptation of Human CD4+ and CD8+ T-Cells to T-Cell Receptor-Mediated Stimulation. Front. Immunol. 8, 1516 (2017).

17. van der Windt, G. J. W. et al. Mitochondrial respiratory capacity is a critical regulator of CD8+ T cell memory development. Immunity 36, 68–78 (2012).

18. Jiang, Y. et al. How synonymous mutations alter enzyme structure and function over long timescales. Nat. Chem. (2022) doi:10.1038/s41557-022-01091-z.

19. Earnest-Noble, L. B. et al. Two isoleucyl tRNAs that decode synonymous codons divergently regulate breast cancer metastatic growth by controlling translation of proliferation-regulating genes. Nat Cancer 3, 1484–1497 (2022).

20. Goodarzi, H. et al. Modulated Expression of Specific tRNAs Drives Gene Expression and Cancer Progression. Cell 165, 1416–1427 (2016).

21. Shen, X., Song, S., Li, C. & Zhang, J. Synonymous mutations in representative yeast genes are mostly strongly non-neutral. Nature 606, 725–731 (2022).

22. Rogalski, M., Karcher, D. & Bock, R. Superwobbling facilitates translation with reduced tRNA sets. Nat. Struct. Mol. Biol. 15, 192–198 (2008).

23. Soto, I. et al. Balanced mitochondrial and cytosolic translatomes underlie the biogenesis of human respiratory complexes. Preprint at https://doi.org/10.1101/2021.05.31.446345.

24. Laricchia, K. M. et al. Mitochondrial DNA variation across 56,434 individuals in gnomAD. Genome Res. (2022) doi:10.1101/gr.276013.121.

25. Lake, N. J. et al. Quantifying constraint in human mitochondrial DNA. bioRxiv 2022.12.16.520778 (2022) doi:10.1101/2022.12.16.520778.

26. Weissensteiner, H. et al. HaploGrep 2: mitochondrial haplogroup classification in the era of high-throughput sequencing. Nucleic Acids Res. 44, W58–63 (2016).

27. van Oven, M. PhyloTree Build 17: Growing the human mitochondrial DNA tree. Forensic Science International: Genetics Supplement Series 5, e392–e394 (2015).

28. Pollard, K. S., Hubisz, M. J., Rosenbloom, K. R. & Siepel, A. Detection of nonneutral substitution rates on mammalian phylogenies. Genome Res. 20, 110–121 (2010).

29. Yonova-Doing, E. et al. An atlas of mitochondrial DNA genotype-phenotype associations in the UK Biobank. Nat. Genet. 53, 982–993 (2021).

30. Jiang, Y. et al. How synonymous mutations alter enzyme structure and function over long time scales. bioRxiv 2021.08.18.456802 (2022) doi:10.1101/2021.08.18.456802.

31. Sookoian, S. & Pirola, C. J. Alanine and aspartate aminotransferase and glutamine-cycling pathway: their roles in pathogenesis of metabolic syndrome. World J. Gastroenterol. 18, 3775–3781 (2012).

32. Gillen, S. L., Waldron, J. A. & Bushell, M. Codon optimality in cancer. Oncogene 40, 6309–6320 (2021).

33. Suzuki, T. et al. Complete chemical structures of human mitochondrial tRNAs. Nat. Commun. 11, 4269 (2020).

34. Gorelick, A. N. et al. Respiratory complex and tissue lineage drive recurrent mutations in tumour mtDNA. Nat Metab (2021) doi:10.1038/s42255-021-00378-8.

35. Martínez-Reyes, I. et al. Mitochondrial ubiquinol oxidation is necessary for tumour growth. Nature 585, 288–292 (2020).

36. Maclaine, K. D., Stebbings, K. A., Llano, D. A. & Havird, J. C. The mtDNA mutation spectrum in the PolG mutator mouse reveals germline and somatic selection. BMC Genom Data 22, 52 (2021).

37. Sanchez-Contreras, M. et al. Multi-tissue landscape of somatic mtDNA mutations indicates tissue specific accumulation and removal in aging. bioRxiv 2022.08.30.505884 (2022) doi:10.1101/2022.08.30.505884.

38. Chen, M. L. et al. Erythroid dysplasia, megaloblastic anemia, and impaired lymphopoiesis arising from mitochondrial dysfunction. Blood 114, 4045–4053 (2009).

39. Lee-Six, H. et al. The landscape of somatic mutation in normal colorectal epithelial cells. Nature 574, 532–537 (2019).

40. Yoshida, K. et al. Tobacco smoking and somatic mutations in human bronchial epithelium. Nature 578, 266–272 (2020).

41. Lee-Six, H. et al. Population dynamics of normal human blood inferred from somatic mutations. Nature 561, 473–478 (2018).

42. Brunner, S. F. et al. Somatic mutations and clonal dynamics in healthy and cirrhotic human liver. Nature 574, 538–542 (2019).

43. Crimi, M., et al. Mitochondrial-DNA nucleotides G4298A and T10010C as pathogenic mutations: the confirmation in two new cases. Mitochondrion 3, 279–283 (2004).

44. Dhindsa, R. S. et al. A minimal role for synonymous variation in human disease. bioRxiv 2022.07.13.499964 (2022) doi:10.1101/2022.07.13.499964.

45. Shen, X., Song, S., Li, C. & Zhang, J. On the fitness effects and disease relevance of synonymous mutations. bioRxiv 2022.08.22.504687 (2022) doi:10.1101/2022.08.22.504687.

46. Kruglyak, L. et al. No evidence that synonymous mutations in yeast genes are mostly deleterious. Preprint at https://doi.org/10.1101/2022.07.14.500130.

47. Sharp, P. M., Bailes, E., Grocock, R. J., Peden, J. F. & Sockett, R. E. Variation in the strength of selected codon usage bias among bacteria. Nucleic Acids Res. 33, 1141–1153 (2005).

48. Boguszewska, K., Szewczuk, M., Kaźmierczak-Barańska, J. & Karwowski, B. T. The Similarities between Human Mitochondria and Bacteria in the Context of Structure, Genome, and Base Excision Repair System. Molecules 25, (2020).

49. Stuart, T., Srivastava, A., Madad, S., Lareau, C. A. & Satija, R. Single-cell chromatin state analysis with Signac. Nat. Methods 18, 1333–1341 (2021).

50. Stuart, T. et al. Comprehensive Integration of Single-Cell Data. Cell 177, 1888–1902.e21 (2019).

51. Hao, Y. et al. Dictionary learning for integrative, multimodal, and scalable single-cell analysis. bioRxiv 2022.02.24.481684 (2022) doi:10.1101/2022.02.24.481684.

52. Hao, Y. et al. Integrated analysis of multimodal single-cell data. Cell 184, 3573–3587.e29 (2021).

53. Garrison, E. & Marth, G. Haplotype-based variant detection from short-read sequencing. arXiv [q-bio.GN] (2012).

54. Li, S. H.-J., Nofal, M., Parsons, L. R., Rabinowitz, J. D. & Gitai, Z. Monitoring mammalian mitochondrial translation with MitoRiboSeq. Nat. Protoc. 16, 2802–2825 (2021).

55. Basak, A. et al. Control of human hemoglobin switching by LIN28B-mediated regulation of BCL11A translation. Nat. Genet. 52, 138–145 (2020).

56. Wang, Q. et al. Rare variant contribution to human disease in 281,104 UK Biobank exomes. Nature 597, 527–532 (2021).

57. Cingolani, P. et al. A program for annotating and predicting the effects of single nucleotide polymorphisms, SnpEff: SNPs in the genome of Drosophila melanogaster strain w1118; iso-2; iso-3. Fly 6, 80–92 (2012).

58. Cunningham, F., et al. Ensembl 2022. Nucleic Acids Res. 50, D988–D995 (2022).

59. Jun, G. et al. Detecting and estimating contamination of human DNA samples in sequencing and array-based genotype data. Am. J. Hum. Genet. 91, 839–848 (2012).

60. Pedersen, B. S. & Quinlan, A. R. Who’s Who? Detecting and Resolving Sample Anomalies in Human DNA Sequencing Studies with Peddy. Am. J. Hum. Genet. 100, 406–413 (2017).

